# CCycDB: an integrative knowledgebase to fingerprint microbially mediated carbon cycling processes

**DOI:** 10.64898/2026.01.28.702190

**Authors:** Jiayin Zhou, Lu Qian, Mengzhi Ji, Yan Li, Kai Ma, Xiaoli Yu, Jiyu Chen, Lu Lin, Xiaofan Gong, Zhili He, Jianjun Wang, Qichao Tu

**Affiliations:** Institute of Marine Science and Technology, Shandong University, Qingdao, China; Southern Marine Science and Engineering Guangdong Laboratory (Zhuhai), Zhuhai, China; State Key Laboratory of Lake and Watershed Science for Water Security, Nanjing Institute of Geography and Limnology, Chinese Academy of Sciences, Nanjing, China; Shandong Key Laboratory of Intelligent Marine Engineering Geology, Environment and Equipment, Qingdao, China

**Keywords:** microbially mediated carbon cycling, functional gene database, shotgun metagenomes, functional composition, taxonomic composition

## Abstract

Microorganisms play essential roles in mediating biogeochemical cycling of carbon across Earth’s ecosystems. Understanding the processes and underlying mechanisms for microbially mediated carbon cycling is therefore critical for advancing global ecology and climate change research. To comprehensively depict these complex biogeochemical processes, we developed CCycDB, a knowledge-based functional gene database, to accurately fingerprint microbially-mediated carbon cycling pathways and gene families, particularly from shotgun metagenomes. The CCycDB database comprises 4,676 gene families classified into six major functional categories, further structured into 45 level-1 and 188 level-2 sub-categories, encompassing a total of 10,991,724 high-quality reference sequences. Validation using both synthetic and real-world datasets demonstrated that CCycDB outperforms existing orthology databases in terms of accuracy, coverage and specificity. By directly targeting carbon-cycling functional gene families, CCycDB provided promising routines to reconstruct both functional gene and taxonomic profiles associated with microbially mediated carbon cycling. Application of CCycDB to shotgun metagenomes from diverse and complex ecosystems revealed pronounced habitat-specific differences in carbon cycling processes and their associated microbial taxa. Collectively, CCycDB provides a powerful and reliable tool for profiling carbon cycling processes from both functional and taxonomic perspectives in complex ecosystems. CCycDB is accessible at https://ccycdb.github.io/.

**Impact Statement:** The microbially mediated carbon cycling processes are the most complex biogeochemical processes in the Earth’s biosphere, playing profound regulatory roles on global climate changes. A key bottleneck in linking microbial communities to global change is the lack of integrated tools for comprehensive carbon cycle profiling. Here, we present CCycDB, a tool that serves a dual purpose—first being a reference database that obtains functional gene and taxonomic profiles and functioning as a customized routine for efficiently aligning sequences and querying associated functional information. CCycDB enables researchers to accurately link microbial community dynamics to carbon cycling and transforming pathways, thereby advancing integrated global change studies with microbes and ecological research via complex metagenomic datasets.

## Introduction

Carbon (C) is the most fundamental element for life, and constitutes the backbone for almost all organic compounds across biospheres of the Earth. It exists in various forms in nature, such as greenhouse gases like carbon dioxide (CO_2_) and methane (CH_4_), as well as complex organic and inorganic compounds. These different forms of carbon compounds are cycled among the atmosphere, land, oceans, and living organisms through various processes (1). C can either be rapidly released to the atmosphere through processes like respiration in the short term (2), or be transformed into long-term recalcitrant C in soils (3) or the vast and deep ocean (4, 5), the latter of which results in C sequestration for geological periods. The form and fate of converted C compounds are expected to have great impacts on the Earth ecosystem, such as global climate change (6).

As the most diverse and abundant life forms on Earth, microorganisms play vital roles are vital drivers of the biogeochemical cycles of various elements (7, 8), including the particularly the C cycle (9, 10). Recent studies have emphasized the importance of microbial C cycling processes in global climate change (11, 12), especially the microbial carbon pump (MCP) processes (13). Multiple studies have even urged to incorporate microbial C processes into ecological models to explain and predict global climate changes (14–16). In the global ocean, there exists a gigantic pool of recalcitrant dissolved organic carbon (RDOC), with an amount equivalent to the atmospheric CO_2_ reservoir (17, 18). The MCP theory provides a compelling explanation for the enigmatic connection between microbial activities and the generation of RDOC, demonstrating that microbial communities are the key players in the long-term sequestration of atmospheric C by the ocean and regulators of global climate change (13).

Since the first industrial revolution in the 18^th^ century, global climate change mainly caused by human activities is rapidly altering almost all the chemical, physical, and biological properties that affect microbial communities (2). Extensive efforts have been made to understand the impacts of climate change on microbial communities and their responsive mechanisms in different ecosystems, especially the microbially mediated C cycling processes (8, 19–21). It has been shown that continuous climate warming and changes in the pattern of temperature fluctuations are expected to accelerate microbial growth and metabolic rates (19, 22–24), further stimulating CO_2_ release, oxygen depletion, and soil C loss (25). In the ocean, CO_2_ accumulated in the atmosphere enters the surface ocean, disrupts the “seawater carbonate buffer system”, causes ocean acidification (6, 26) and shallows the “surface ocean mixed layer” (2), thereby affecting the potential of ocean to act as C sinks (4, 27). Therefore, disentangling the microbial processes, especially C cycling processes is of critical importance to better understand the roles that microbial communities play and their corresponding feedbacks in the Earth’s ecosystems.

The rapid development of high-throughput metagenomic sequencing technologies that provide community-level information (28) has made it a routine practice to investigate the complex microbial communities in various ecosystems. In utilizing these approaches, a persistent challenge is to accurately and efficiently mine biological/ecological information (e.g., taxonomic and functional gene information) from massive amounts of sequencing data.

Over the past decade, large public orthology databases, such as Clusters of Orthologous Groups (COG) (29), evolutionary genealogy of genes -- Non-supervised Orthologous Groups (eggNOG) (30), and Kyoto Encyclopedia of Genes and Genomes (KEGG) (31), have been widely used in metagenomic studies for their comprehensive information and potential to interpretate the datasets. However, the massive amount of information generated by these databases makes it difficult for researchers to accurately and efficiently obtain useful and specific information. Recently, small-scale databases with high coverage and accuracy have been developed to profile certain metabolic pathways, such as Carbohydrate-Active Enzymes (CAZy) for carbohydrate-active enzymes (32), CARD for antibiotic resistance genes (33), NCycDB for nitrogen cycle (34), SCycDB for sulfur cycle (35) and PCycDB for phosphorus cycle (36). These specific knowledgebases have greatly facilitated researchers to accurately and efficiently capture the information of targeted gene families. However, to our best knowledge, a comprehensive and highly accurate functional gene database specifically targeting microbially mediated C cycling processes remains urged, even though carbon has become “a matter of public concern” as a result of global climate change.

In this study, a knowledge-based functional gene database termed CCycDB (https://ccycdb.github.io/) was developed to profile microbially mediated C cycling processes in complex environments. CCycDB represented the most up-to-date knowledge of gene families involved in the carbon cycle with 4,676 gene families classified into six categories, which were further categorized into 45 level-1 and 188 level-2 sub-categories. Together with the database, a metagenomic profiling pipeline was developed to recover microbial gene families and taxonomic groups involved in C cycling processes from both reads and assembled contigs. Gene families and their homologs from public databases were integrated with CCycDB to improve the accuracy and reduce false positives. We applied CCycDB to analyze both simulated and real-world metagenomic datasets, and demonstrated its superior performance comparing to other large public orthology databases. Collectively, CCycDB is a useful and powerful tool to fingerprint microbially mediated carbon cycling gene families and processes in complex ecosystems.

## Methods

### CCycDB framework development and annotation overview

An integrative and sophisticated pipeline was developed to construct CCycDB (Figure 1), which was based on our previous framework in developing NCycDB (34) but with moderate modifications, such as incorporating multiple knowledgebases (e.g., CAZy and BRENDA) and supporting multiple data input (e.g., both contigs and raw data). The pipeline encompasses procedures including databases construction, performance assessment, and metagenomic profiling.

**Figure 1.**
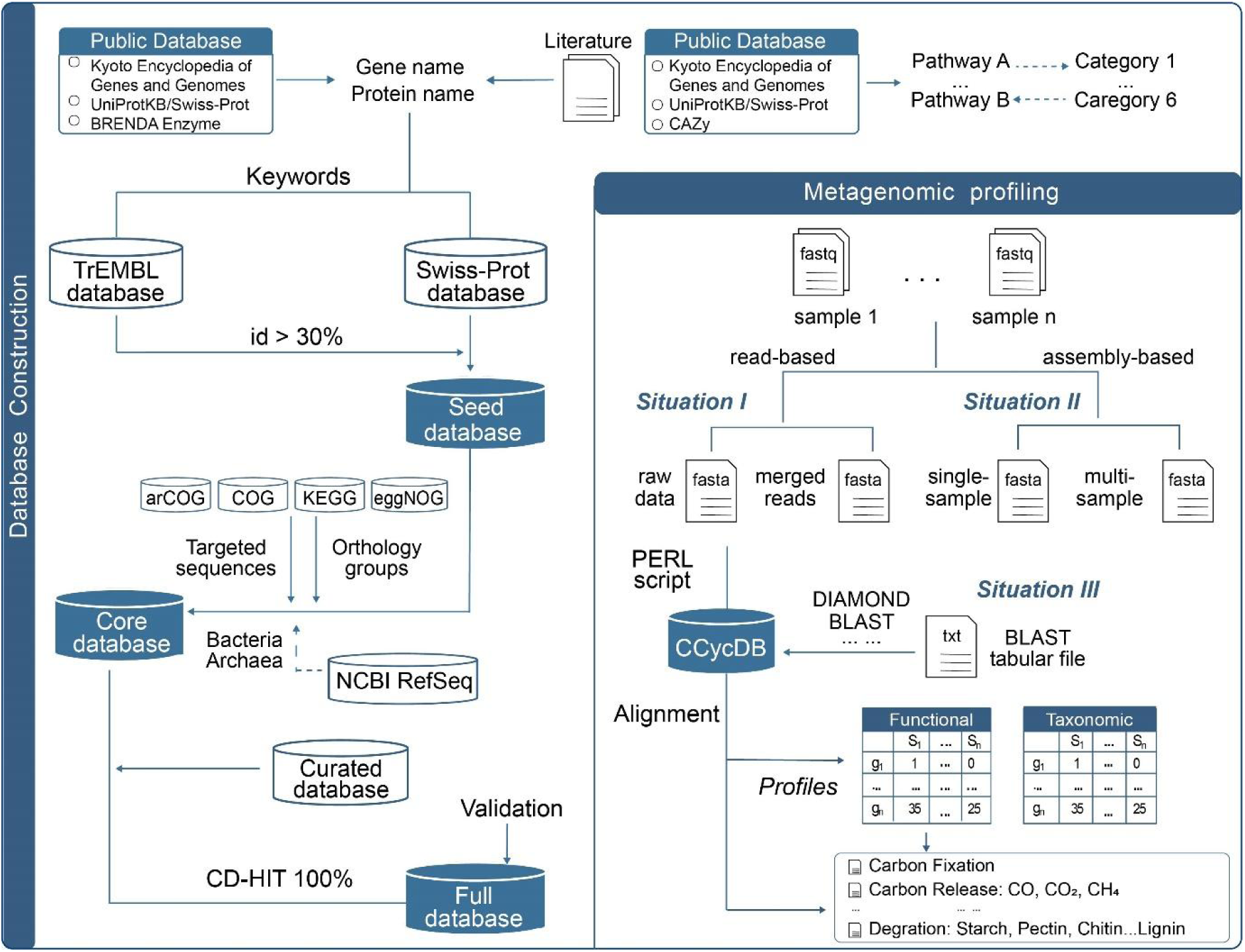
Workflow summary for constructing CCycDB. Five major components were included in this framework, including seed database construction, core database construction, full database construction, performance assessment and metagenomic profiling. Metagenomic profiling supports “read-based” and “assembly-based” data processing approaches of shotgun metagenomes to generate functional and taxonomic profiling.

**Figure 2.**
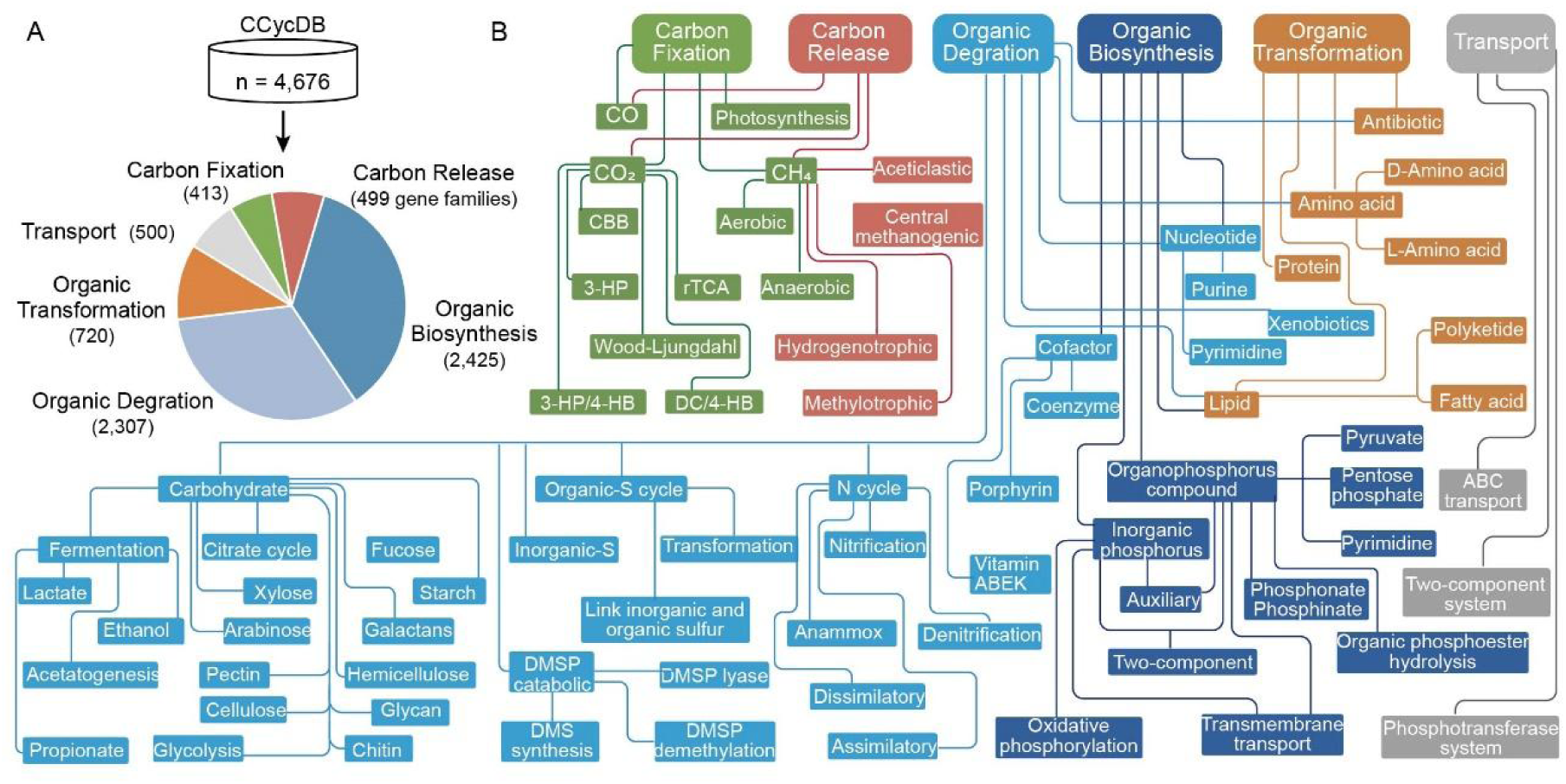
Summary of gene families and categories in CCycDB. (A) A total of 4,676 gene families were grouped into six categories based on the source and fate of carbon, including carbon fixation, carbon release, organic biosynthesis, organic degradation, organic transformation, and transporters. (B) Flow charts shows the classification in CCycDB. Each category contains a series of different specific pathways. Different colors represent different category.

### Database sources

Keywords searching and sequence retrieval for the seed database were conducted at the UniProt website (https://www.uniprot.org/) (37). Enzyme information obtained from Braunschweig Enzyme Database (BRENDA) (https://www.brenda-enzymes.org/) was used to inspect and correct the annotation. Multiple orthologous databases were integrated and used for homologous gene identification in this study, including Archaeal Clusters of Orthologous Genes (arCOG) (https://ftp.ncbi.nlm.nih.gov/pub/wolf/COGs/arCOG/), Clusters of Orthologous Groups of proteins (COG) (https://ftp.ncbi.nlm.nih.gov/pub/COG/), Kyoto Encyclopedia of Genes and Genome (KEGG) (http://www.genome.jp/kegg/) and evolutionary genealogy of genes: Nonsupervised Orthologous Groups (eggNOG) (http://eggnogdb.embl.de/download/eggnog5.0/). The NCBI non-redundant RefSeq databases of archaea and bacteria were downloaded from https://www.ncbi.nlm.nih.gov/refse/release.

Curated small-scale databases (SCycDB, MCycDB, PCycDB, and VB_12_Path) were downloaded from https://github.com/qichao1984 and https://github.com/ZengJiaxiong.

### Seed database construction

High-quality reference sequences for the targeted gene families were collected for the seed database construction. The microbially mediated carbon cycling encompasses many sophisticated processes and is the most complex element cycle on Earth, making it challenging to determine the gene families involved in each process. By referring to multiple literatures, KEGG pathway/module (31), CAZy (32) and its associated tools (e.g. dbCAN2 (38)), DRAM (Distilled and Refined Annotation of Metabolism) (39) and the previously developed curated functional gene databases such as SCycDB (35), PCycDB (36), MCycDB (40) and VB_12_Path (41), a number of carbon cycling pathways and corresponding gene families and high quality sequences were recruited in CCycDB. Based on the substrates, functional processes and/or the fate of C metabolism, the targeted gene families were grouped into six different categories, including carbon fixation, carbon release, organic biosynthesis, organic degradation, organic transformation and transporters. For each of these categories, many different sub-categories and pathways were covered. Manual inspection revealed that gene families performing the same function were sometimes inconsistently tagged in different databases. Thus, to ensure the accuracy of functional gene annotations, the gene family names were manually inspected and corrected based on literatures and public databases (e.g., KEGG, Swiss-Prot (42) and BRENDA (43)). Then, keywords were generated based on one-to-one correspondence between gene families and corresponding protein names. Sequences were collected from the Swiss-Prot database (42) and the TrEMBL database based on curated keywords to form the seed database.

### Core and full database construction

We recruited four publicly available orthology databases, including arCOG (44), COG (29), eggNOG (30), and KEGG (31), and searched against them the seed database using USEARCH (45) with 70% identity. Homologous gene families were also identified to diminish false-positive effects due to “small database issue” (34). The NCBI archaeal and bacterial RefSeq databases were retrieved and merged, greatly improving the integrality of CCycDB. Considering the completeness and accuracy of targeted functional gene database sequences, we further analyzed and and integrated several manually curated small-scale databases, including SCycDB (35), PCycDB (36), MCycDB (40) and VB_12_Path (41), to complement the coupling relationship with sulfur cycle, methane cycle, phosphorus cycle and B_12_ biosynthesis pathways, respectively. Finally, duplicated sequences were removed using the CD-HIT program (46) at 100% identity cutoff. The nonredundant representative sequences and homolog sequences were retained to construct the CCycDB.

### Functional and taxonomic profiling in CCycDB

Both functional and taxonomic profiling were implemented in CCycDB. Two common data processing approaches were supported, including read-based (merged reads or raw data) and assembly-based (single-sample or multi-sample assembly) (Figure 1). Users can specify the “-situation” parameter to use read files or assembly-derived files as input. For the assembly-based profiling, users are required to utilize assembly software (e.g., MEGAHIT (47), and metaSPAdes (48)) beforehand to obtain contigs. Prediction of open reading frames (ORFs) of contigs can be done with public software (e.g., Prodigal (49), and FragZGeneScan (50)). The predicted coding sequences can be used as input files for the assembly-based mode. The program “CCycDB.PL” calls a user-specified searching tool (e.g., DIAMOND, BLAST, USEARCH; default option: -e 1e-5 -id 30) to generate the best-hit mapping. Furthermore, if the input files are tabular files (-situation tabular) generated by searching tool, the alignment step will be omitted. Based on the alignment results, SEQ2GENE (-situation read-based or -situation tabular) or ORF2GENE (-situation assembly-based) provides the corresponding gene information for targeted sequences. Finally, the results are presented as a tab-delimited abundance table. Users can further specify the -norm parameter to normalize the abundance table. If -norm 0 is chosen, random subsampling will not be executed. If -norm 1 is chosen without specifying -rs (number of sequences for random subsampling), all samples will be normalized to the lowest number of sequences by default. In addition, descriptions of gene families from other public databases are also provided to facilitate data comparison and analysis.

### Benchmarking CCycDB against public databases

We benchmarked the performance of CCycDB in terms of coverage, specificity and accuracy. Two types of datasets were used, including (i) 25 real shotgun metagenomic sequencing datasets derived from seven distinct habitats, and (ii) an artificial dataset constructed from the NCBI GenBank database. The artificial dataset included a total of 69,590 sequences belonging to 755 genes from the NCBI GeneBank database with 53,460 related sequences of 707 carbon metabolism genes and 16,130 unrelated sequences of 48 other gene families. This artificial dataset was searched against the CCycDB and KEGG using the DIAMOND program (option: -k 1 -e 1e-5 -id 30), and against eggNOG using eggNOG-mapper (-m diamond -evalue 1e-5) (51). The annotation performance was evaluated by five indices (recall, specificity, precision, accuracy and F1-score) defined below:

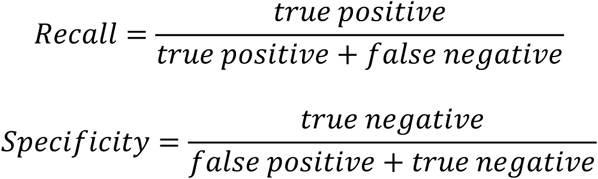

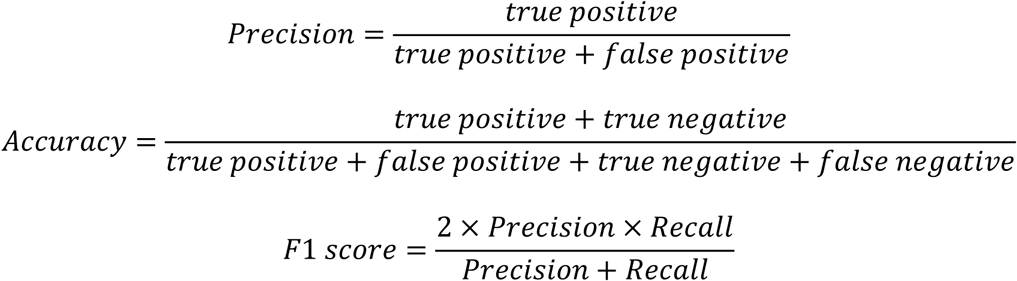

where false positives included the relevant sequences that were misassigned or not correctly annotated and the unrelated sequences that were misassigned to C cycling gene families, and false negatives were the relevant sequences that were not assigned to the corresponding C cycling gene families. Only the sequences that were correctly assigned and annotated were considered as true positives.

### Case study

To evaluate the performance of CCycDB in profiling complex metagenomes, we applied CCycDB to analyze microbially mediated C cycling processes and taxonomic groups for 25 real-world shotgun metagenomic datasets derived from seven distinct habitats, including the oceanic surface water layer (SRF, n=4), deep chlorophyll maximum layer (DCM, n=3), mesopelagic zone (MES, n=3), aquaculture (n=4), forest soil (n=3), mangrove sediments (n=4), and permafrost (n=4). The metagenomic datasets were assembled using MEGAHIT (version 1.2.9, options: --k-min 39 --k-max 109) (47). Gene prediction was performed on resulting contigs > 500bp using FragGeneScan (version 1.31, options: -complete=1 -train=complete) (50). The predicted coding sequences were subsequently searched against CCycDB with the DIAMOND program (option: -k 1 -e 0.00001 -id 0.3) (52). Predicted coding sequences belonging to targeted functional gene families were extracted using the seqtk program (53). Bowtie (version 2.4.4) (54) was used to align extracted sequences to raw reads. PERL script was then used to generate functional profiles. The Kraken2 program (55) was used to perform taxonomic assignments of the extracted sequences. Then, the generated functional and taxonomic profiles were applied for subsequent statistical analyses.

## Results

### Summary of microbially mediated C cycling gene families in CCycDB

A total of 10,991,724 high-quality reference sequences covering 4,676 gene families were recruited for CCycDB, targeting a variety of microbially mediated C cycling processes. Furthermore, a total of 3,219,403 homologous sequences were also included in CCycDB to minimize false positive assignments resulting from small database issues (34). Of the homologous sequences, 53,381 were from the arCOG, 326,342 from the COG, 1,642,871 from the eggNOG, and 1,257,245 from the KEGG database (Table 1). CCycDB is categorized by six categories, including carbon fixation, carbon release, organic biosynthesis, organic degradation, organic transformation, and transporters. For each category, a series of sub-categories and pathways were defined based on the involved substrates (Table 2). Notably, as the most complex biogeochemical cycle in the Earth’s biosphere, sharing of gene families and processes by different pathways is a common scenario. A brief description of CCycDB was provided below with more details in Table 1, Table 2 and Supplementary Table 1.

### Carbon fixation

Microbial functional gene families that convert simple gaseous carbon into more complex organic carbon were classified as carbon fixation. This category comprised four level-1 sub-categories, 17 level-2 sub-categories and 21 pathways. Microbially mediated carbon fixation processes for three major types of gaseous carbon were characterized, including carbon monoxide (CO), carbon dioxide (CO_2_), and methane (CH_4_). For microbial fixation of CO, carbon monoxide dehydrogenase (CODH) that catalyzes the key reaction of CO oxidation to CO_2_, as well as gene families that mediate reactions involving CO were included (56, 57). For CO_2_ fixation, in addition to the most widespread Calvin-Benson-Bassham (CBB) cycle on Earth, gene families involved in five other CO_2_ fixation pathways were also covered, including the reductive tricarboxylic acid cycle (rTCA), reductive acetyl-CoA (WL), 3-hydroxypropionate bicycle (3-HP), 3-hydroxypropionate/4hydroxybutyrate (3-HP/4-HB) and dicarboxylate/4-hydroxybutyrate cycles (DC/4-HB). For CH_4_ fixation, gene families responsible for aerobic and anaerobic oxidation of methane were covered. A total of 413 gene families were included for carbon fixation, of which 16 gene families are involved in CO fixation, 128 for CO_2_ fixation, and 207 for CH_4_ fixation pathways. In total, 1,279,437 targeted sequences and 310,628 homologous sequences were collected for the targeted carbon fixation gene families.

### Carbon release

In addition to carbon fixation, microorganisms also consume organic carbon to obtain energy and release gaseous carbon via processes like respiration. The balance between carbon fixation and release is crucial for global climate change (1). Here, the carbon release category comprised three level-1 sub-categories covering six level-2 sub-categories. Enzymatic reactions directly involved in releasing gaseous carbon (e.g., CO and CO_2_) were mainly referred for recruiting carbon release gene families. For example, isocitrate dehydrogenase encoded by the *icd* gene catalyzes the oxidative decarboxylation of isocitrate, producing α-ketoglutarate and CO_2_ (58). For CH_4_ release, gene families involved in methanogenesis pathways associated with commonly used substrates (e.g., H_2_ or formate, acetate and methylated compounds) were recruited. According to the form of gaseous carbon, the three level-1 sub-categories included CO (23 gene families), CO_2_ (337 gene families) and CH_4_ (181 gene families). A total of 499 gene families were included in this category, covering 1,070,260 sequences and an additional of 278,093 homologous sequences.

### Organic biosynthesis

Like other living organisms, microorganisms also synthesize complex organic compounds from much simpler molecules, via multi-step enzyme-catalyzed processes. Notably, a variety of organic compounds can only be synthesized by microorganisms, such as the production of cobalamin in natural ecosystems (59, 60). For organic biosynthesis, a series of microbial gene families responsible for the biosynthesis of various organic compounds were targeted, including photosynthesis, the biosynthesis of cofactors, carbohydrates, glycan, nucleotides, amino acids, lipid, secondary metabolites, antibiotics, and more. These different sub-categories were further characterized into 85 processes according to the products that microorganisms synthesize. A total of 2,425 gene families and 7,112,337 sequences were recruited for 12 level-1 sub-categories, covering 55 level-2 sub-categories and 85 pathways. In addition, 1,976,321 homologous sequences were identified and included.

### Organic degradation

Functioning as major decomposers in the Earth’s biosphere, microorganisms break large complex organic compounds into smaller molecules to obtain energy and nutrients, allowing the released elements to reenter the global biogeochemical cycles (61). Recent studies demonstrate that microorganisms also hold the genetic potential to degrade almost all kinds of organic compounds on Earth, including complex human-made materials such as plastics (62, 63). A total of 15 level-1 sub-categories were characterized for organic degradation based on substrates, including amino acids, antibiotics, aromatics, carbohydrates, cofactors, lipids, methane, nucleotides, nitrogen, organic and inorganic sulfur, DMSP, secondary metabolites, xenobiotics and others. Among these, the carbohydrate category encompassed key processes such as the citrate cycle, glyoxylate cycle and oxidative phosphorylation, and substrate-levels based on the degree of lability or recalcification (e.g., starch, hemicellulose, cellulose, pectin). Since microorganisms cannot synthesize lignin, lignin degradation was characterized within the xenobiotic degradation sub-category. A total of 2,307 gene families with 5,677,874 sequences and 1,832,148 homologous sequences were recruited for organic degradation (Table 1).

### Organic transformation

Organic transformation is defined as the process that complex organic substances are transformed into other compounds through various biochemical reactions or metabolic pathways. Here, organic compounds less degradable by microorganisms were mainly focused. For example, most D-amino acids are difficult to be used by bacteria as a single carbon or nitrogen source, and they are considered as recalcitrant dissolved organic matter (64, 65). A total of 720 gene families were recruited for organic transformation, in which 1,113,217 sequences and 408,355 homologous sequences were present. These gene families were further divided into seven level-1 sub-categories that cover 15 level-2 sub-categories and 19 pathways, including organic transformation of proteins (13 gene families), carbohydrates (8 gene families), amino acids (19 gene families), glycan (139 gene families), lipid (214 gene families), antibiotics (147 gene families) and xenobiotics (248 gene families).

### Transporters

Besides the abovementioned categories, a total of 500 gene families involved in phosphotransferase systems, ABC transport systems, two-component systems and other transport systems were also recruited, with a total of 1,341,038 sequences and 549,764 homologous sequences included in the database.

### Taxonomic composition of carbon cycling pathways in CCycDB

During the construction of the database, currently sequenced microbial genomes from NCBI RefSeq database were also integrated to improve the comprehensiveness of CCycDB. The sequenced microorganisms with genes mapped to CCycDB were extracted to further explore the taxonomic information of microorganisms potentially participated in different carbon cycling pathways. Consistent with the current sequencing efforts, bacterially originated sequences comprise around 99.30% of all sequences in CCycDB with a total of 48 phyla, 82 class, 197 order, 463 families, 2,622 genera and 23,383 species in CCycDB (Table 3 and Supplementary Figure 1). Only 0.70% of the sequences were detected to be derived from archaeal genomes with a total of six phyla, 12 class, 23 order, 41 families, 142 genera and 591 species (Table 3).

### CCycDB is more comprehensive and accurate than other public orthology databases

The performance of CCycDB was evaluated by comparing it with other commonly used publicly available orthology databases from multiple angles, such as accuracy, coverage and specificity. First, the coverage of carbon cycling gene families in CCycDB far exceeded other large databases. A total of 4,676 gene families were covered in CCycDB, while only 529, 1607, 2266 and 4184 gene families were found in arCOG, COG, eggNOG and KEGG, respectively (Figure 3A). Multiple functionally important gene families involved in different carbon cycling pathways were present in CCycDB but absent (or not clearly annotated) from these three orthology databases. For example, *icdA* for carbon dioxide release, *hcd* for 3-hydroxypropionate/4hydroxybutyrate cycle, *dddKQWY* for organic sulfur transformation, *vhoACG* for methanophenazine hydrogenase and *ppd* (phosphonopyruvate decarboxylase) for organic phosphorus metabolism were absent from these public databases. At the meanwhile, CCycDB also provided more detailed information at the substrate and pathway levels. Second, CCycDB was highly accurate regarding the definition of gene families. A common issue existing in public orthology databases is that multiple gene families are always mixed together as one orthologous group due to high homology among different gene families (e.g., *pmoA* and *amoA*, *psrA* and *phsA*) (34).To avoid such situations, all gene families included in CCycDB were carefully distinguished and manually inspected, rather than the conventional generation of orthologous groups by unsupervised clustering approaches. Third, the accuracy and specificity of CCycDB was further validated by comparing with KEGG and eggNOG databases using an artificial dataset containing both related and unrelated sequences. The performance was quantified using five commonly used metrics, including recall, specificity, precision, accuracy and F1-score. The results indicated that CCycDB outperformed KEGG and eggNOG databases in all of these metrics. CCycDB held the advantage in distinguishing homologous gene families during database searching, resulting in a much lower false-positive rate and overall higher accuracy than other databases. Specifically, CCycDB achieved an overall accuracy of 99.6% and F1-score of 0.99 among the 755 randomly selected gene families. In contrast, KEGG and eggNOG databases respectively achieved 72.84% and 84.10% accuracy, and 0.78 and 0.89 F1-score (Figure 3B). High false-positive rates were observed for public orthology databases. CCycDB was characterized with an overall 99.51% recall, 99.97% specificity, 99.99% precision and low false-positive rate of 0.02%, while KEGG and eggNOG had an overall 66.27% and 84.19% recall, 94.64% and 94.51% specificity, 97.61% and 94.51% precision, 5.36% and 16.18% false-positive rates, respectively (Figure 3 and Supplementary Table 3).

**Figure 3.**
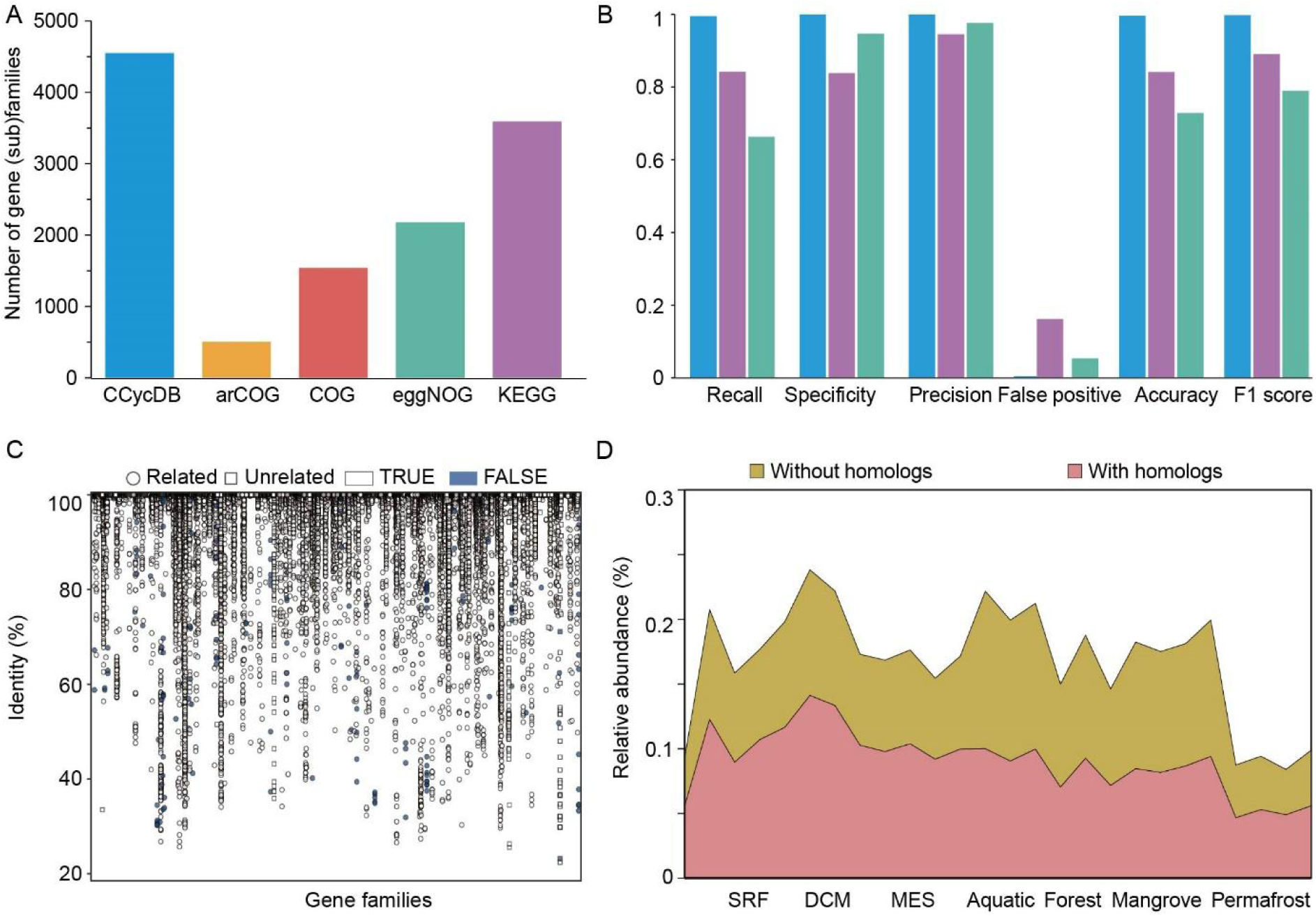
Performance and validation of CCycDB in terms of coverage, specificity and accuracy. (A) The number of gene families involved in carbon cycling processes detected in different databases. (B) Accuracy of different databases validated by an artificial dataset. Different colors represent different databases. Across 755 randomly selected gene families, CCycDB achieved 99.6% overall accuracy and 0.99 F1-score, outperforming KEGG (72.84% accuracy, 0.78 F1-score) and eggNOG (84.10% accuracy, 0.89 F1-score). For additional metrics, CCycDB exhibited 99.51% recall, 99.97% specificity, 99.99% precision, and a low false-positive rate of 0.02%. In contrast, KEGG and eggNOG showed lower recall (66.27% and 84.19%), specificity (94.64% and 94.51%), precision (97.61% and 94.51%), and higher false-positive rates (5.36% and 16.18%), respectively. (C) Validation of the accuracy of sequences annotated by CCycDB using an artificial dataset. (D) Illustration of the “small database issue” via real shotgun metagenomic datasets. The results showed that about 53.69% of sequences were better mapped to homologs than targeted gene families.

### Application of CCycDB to profile complex environmental samples

Shotgun metagenomic sequencing datasets from seven distinct habitats, including the oceanic surface water layer, deep chlorophyll maximum layer, mesopelagic zone, aquaculture, forest, mangrove and permafrost, were collected and annotated against CCycDB to profile microbial carbon cycling gene families and taxonomic groups. Detrended correspondence analyses based on Bray-Curtis dissimilarity suggested that both functional and taxonomic compositions were significantly different (*P* < 0.05) among these seven habitats (Figure 4 and Supplementary Figure 2). Notably, more gene families (2700 ± 46) were detected in mangrove sediment samples than those in other habitats, particularly for C fixation (277 ± 4) and C release (305 ± 6) processes. Among these, gene families involved in the WL pathway and anaerobic oxidation of methane were consistently abundant across all samples (Figure 5). The surface ocean and forest samples were found with fewer gene families involved in C cycling (2026 ± 64 and 1944 ± 98, respectively). Specifically, the surface ocean was found with the most sequences involved in carbon fixation via photosynthesis and the fewest sequences related with carbon degradation processes of less labile carbon, such as hemicellulose (0.31%), xylan (0.26%), cellulose (0.23%) and pectin (0.23%) degradation, whereas forest microbial communities were found with the fewest gene families belonging to carbon fixation (11.99%) and carbon release (9.25%) processes (Figure 4, Figure 5 and Supplementary Figure 4). Taxonomically, *Pelagibacteraceae* (30.13%), *Prochloraceae* (24.05%) and *Alteromonadaceae* (11.48%) were predominant microbial families associated with C cycling in ocean samples, while *Enterococcaceae* (74.16%) was the most abundant C cycling in the forest soil, followed by *Bacillaceae* (6.15%), *Streptococcaceae* (3.28%) and *Bradyrhizobiaceae* (2.45%) (Supplementary Figure 2). In the mangrove and permafrost samples, *Mycobacteriaceae* (7.81% and 3.50%), *Bradyrhizobiaceae* (4.58% and 14.49%) and *Streptomycetaceae* (9.05% and 4.33%) were the predominant groups (Supplementary Figure 2). For gaseous carbon fluxes jointly controlled by C fixation and C release, the predominant families associated with carbon fluxes were significantly different (p < 0.05) across different habitats. The predominant groups recovered from ocean included *Pelagibacteraceae*, *Prochloraceae* and *Alteromonadaceae*, while *Vibrionaceae* and *Rhodobacteraceae* were the most abundant groups in the aquaculture system. *Enterococcaceae* and *Bacillaceae* predominated in the forest soil, while *Mycobacteriaceae* and *Rhodobacteraceae* were the most abundant in mangrove sediment (Figure 5 and Supplementary Figure 5). These results demonstrated that both C cycling gene families and their carrying microbial taxa greatly differ among different habitats.

**Figure 4.**
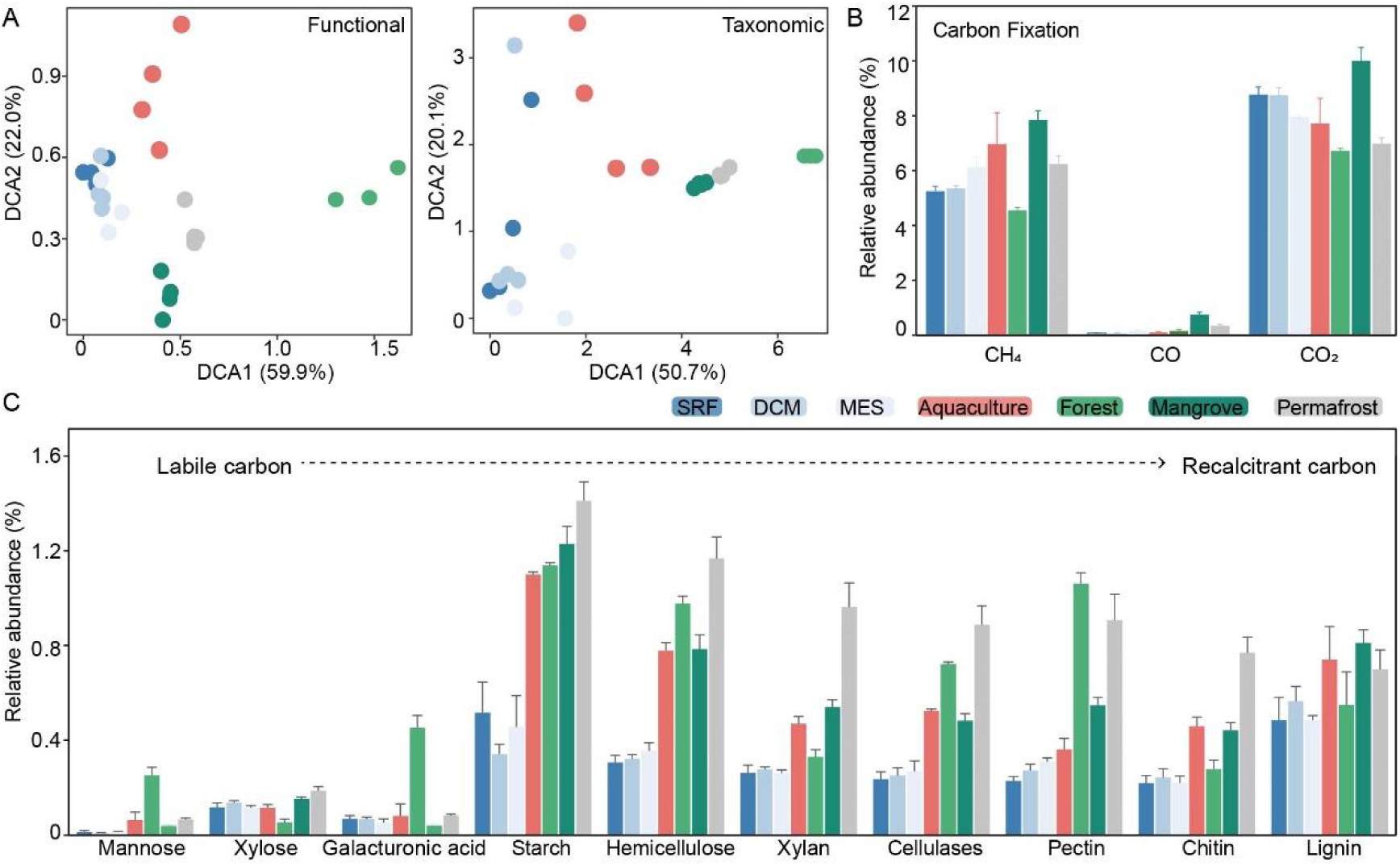
Application of CCycDB to characterize carbon cycling processes in seven distinct habitats. (A) Detrended correspondence analysis of carbon cycling gene families and taxonomy at the species level. A clear separation between different habitats could be observed. Relative abundance of (B) carbon fixation and (C) a subset of organic degradation gene families in different habitats. Different colors represent different habitats. SRF, surface water layer; DCM, deep chlorophyll maximum layer; MES, mesopelagic zone.

**Figure 5.**
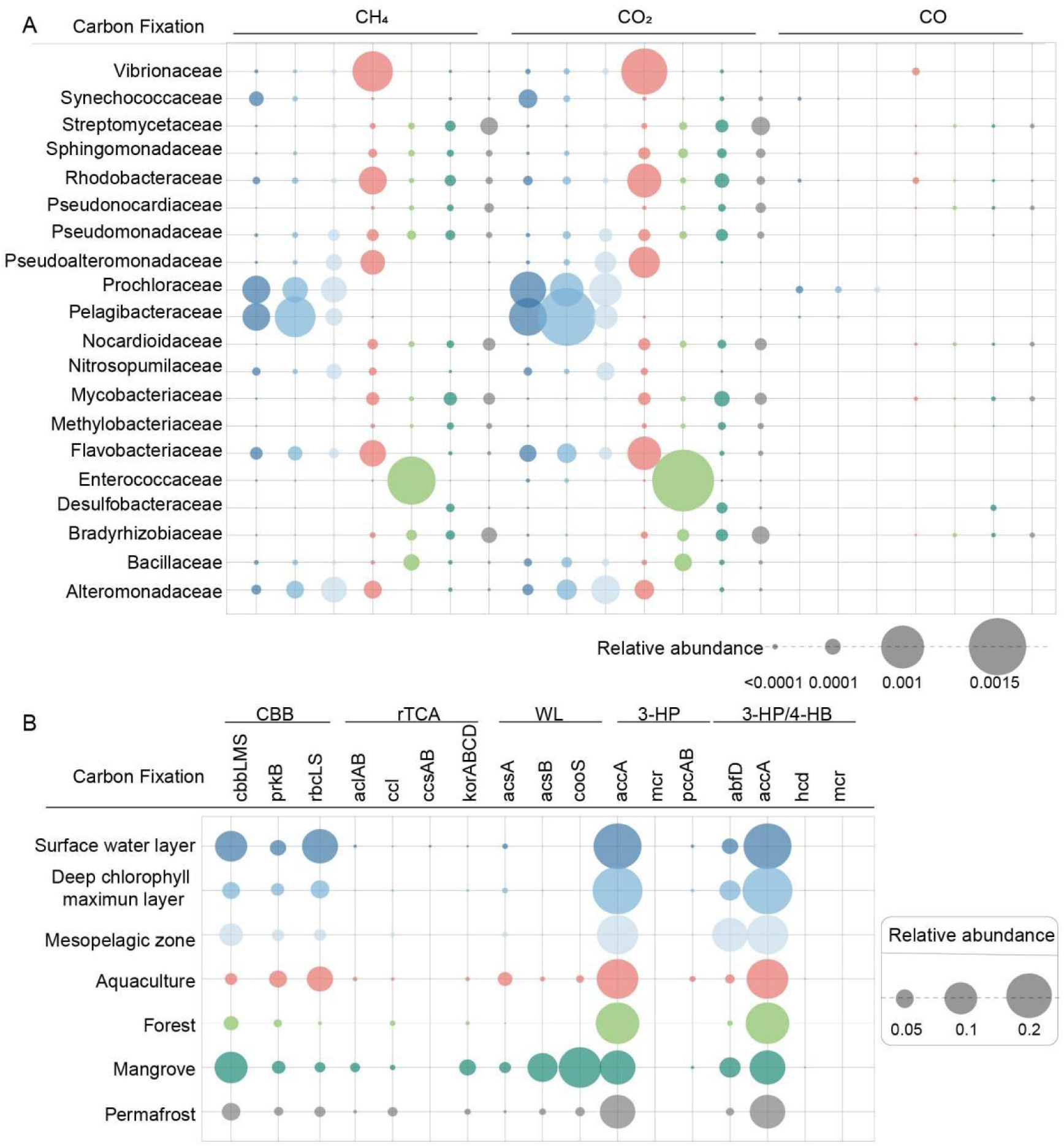
The relative abundance of predominant microbial taxonomic groups (A) and key gene families (B) involved in carbon fixation in different environments.

## Discussion

The C cycle is the most complex and vital biogeochemical cycle in Earth’s ecosystems, and is essential for maintaining ecosystem multifunctioning and stability. Microbial communities are key participants in the C cycling processes, and modulate the direction of global climate change (12, 13, 66). For instance, marine bacteria and archaea are responsible for the respiration of most of the carbon that sinks into the dark ocean (67). Characterization of the structure of typical refractory organic carbon matter (e.g., carboxyl-rich alicyclic molecules) in the deep ocean has revealed their microbial origin (68). Thus, exploring the function and taxonomy of microbial communities involved in various C cycling processes is of critical significance for understanding the ecosystem functionality and stability, especially under global events like climate change.

The developed CCycDB can provide accurate functional and taxonomic profiles of microbially mediated C cycling processes in complex environments. Accuracy, which determines the precision of obtained metagenomic profiles and further statistical results, is the most crucial issue for such knowledge-based functional gene databases. The construction of CCycDB employs a sophisticated knowledge-based database construction procedure (34) that has been extensively validated for its high accuracy. These databases have been now extensively used to understand microbially mediated biogeochemical cycles and underlying mechanisms in various ecosystems via shotgun metagenomic sequencing datasets (69–71). In the database construction procedure, the accuracy of CCycDB is guaranteed both at the gene family and the sequence levels. At the gene family level, a common issue in automatically generated large databases (e.g., arCOG, COG and eggNOG) is the grouping of multiple gene families within one single group. For example, polysulphide reductase (*phsA*) and thiosulphate reductase (*psrA*) involved in the organosulpur cycle, and uroporphyrinogen-III c-methyltransferase (*cobA*) of cobalamine aerobic synthesis and siroheme synthase (*cysG*) of anaerobic processes are always mixed together because of their high homology. The *pmoA* gene family encoding methane monooxygenase and *amoA* gene family encoding ammonia monooxygenase share the same KO identifier (K10944) in KEGG. However, these gene families play different roles in different pathways, and such a mixture of multiple gene families in a single orthologous group may pose challenges for researchers in analyzing these gene families, leading to overestimated gene abundances and ambiguous interpretation of observed results. To mitigate this, gene families were distinguished by establishing one-to-one relationship between gene families and functional annotations when recruiting sequences for seed database construction. At the sequence level, the accuracy of all sequences was further improved by introducing homologous gene families from public orthology databases when constructing the core database. When the real metagenomic sequencing datasets were searched against the database with or without homologs, up to 53.69 ± 1.14% of the sequences showed closer relationship with homologs than the targeted gene families. This suggests that introducing homologous gene families can substantially reduce false positive assignments and improve accuracy. In the testing with the artificial dataset, CCycDB showed superior performance in both recalling positive match and avoid misassignments to unrelated gene families. In addition, we have also noticed that sequences for some genes (e.g., *pccA* and *PCCA*) are derived from eukaryotes rather than prokaryotes, thus the source of collected sequences was reviewed to exclude the eukaryotic portion.

CCycDB is not only a functional gene database (i.e., collection of sequences) with high accuracy but also serves as a knowledgebase with comprehensive gene families and reliable annotation and categorization. Incorporating the most up-to-date knowledge of microbial C cycling processes and genes enables us to comprehensively analyze microbial communities and their relationships with the changing environment. The completeness of CCycDB is reflected both in gene families and pathways involved in carbon cycling. With current effort, CCycDB encompasses 4,676 gene families and 188 level-1 sub-categories with 10,991,724 high-quality sequences involved in six categories. The covered gene families not only include common and key gene families, such as *cbbLS* or *rbcL* encoding forms I ribulose-1,5-bisphosphate carboxylase/oxygenase (RuBisCO) and *cbbM* encoding forms II RuBisCO for Calvin cycle, *acsA* encoding CO dehydrogenase and *acsB* encoding acetyl-CoA synthase for Wood-Ljungdahl (WL) pathway for carbon fixation (72, 73), *dddDKLPQWY* encoding dimethylsulphoniopropionate (DMSP) lyase for DMSP degradation (74–76), and *pmoABC* for the initial oxygenation of methane to methanol in methanotrophs (77, 78). Many gene families that are currently absent from others databases but play important roles in certain pathways are also included. For instance, the gene families *hcd* (encoding 4-hydroxybutyryl-CoA dehydratase) and *mmtN* (encoding methionine methyltransferase), which are a critical enzyme for 3HP/4HB (79, 80) and the marker gene of DMSP, are missing in almost all public orthology databases. Representative sequences of these gene families were collected from public literature (81–83). In addition, CCycDB categorizes functional genes/proteins into specific pathways. For instance, methanogenesis are broadly characterized as hydrogenotrophic (H_2_ or formate as substrate), aceticlastic (acetate), and methylotrophic (methylated compounds) on the basis of substrate use. The newly discovered reversed oxidative tricarboxylic acid (roTCA) cycle of autotrophic CO_2_ fixation is highly efficient, which utilizes citrate synthase instead of ATP-citrate lyase in contrast to the rTCA cycle (84). However, the roTCA cycle can hardly be recognized by bioinformatic prediction (84, 85), thus it is not specifically categorized in CCycDB. In addition, a series of previously developed curated functional gene databases (35, 36, 40, 41) have also been integrated to complement the gene catalogs of CCycDB. Besides that, the CAZy database, an essential resource for studying complex carbohydrate metabolism (32), has also been referenced and further incorporated to complement CCycDB. CAZy identifiers or EC numbers were categorized at substrate levels to enrich the annotation of organic compound degradation category in CCycDB. Glycoside hydrolases (GHs) are involved in catalyzing the hydrolysis of glycosidic bonds of glycosides, while polysaccharide lyases (PLs) and carbohydrate esterases (CEs) are usually associated with the degradation of organic carbon (86). According to the recalcitrance of organic carbon components, the degradation of easily mineralizable starch is commonly attributed to the GH-13/15/31 of CAZy (87). PLs (e.g., PL-1, 3, 4, 9), GHs (GH-28, 78, 88, 95, 105, 115) and CEs (e.g., CE-8, 12) are often related to pectin degradation (88). Enzymes responsible for the decomposition of recalcitrant carbon components, such as chitinase, are listed in GH18 and GH19, with a few chitinases appears in GH23 and GH48 (89, 90). Auxiliary Activities (AAs) are related to lignin decomposition (88, 91). Consequently, CCycDB can be used as a functional gene database for metagenomic profiling, as well as a knowledgebase for better understanding the microbial C cycling processes and underlying mechanisms.

The potential coupling mechanisms of microbially driven C cycle with sulfur, nitrogen and phosphorus cycles have also been critically considered in CCycDB. For example, the dimethylsulfoniopropionate (DMSP) catabolic pathway can be used as a major source of carbon and sulfur for diverse bacteria. Gene families involved in DMSP synthesis (*dsyB* and *mmtN*) and lyase (*dddDKLPQWY*), demethylation (*dmdABCD*), and ancillary (*dddACT*, *acuIKN* and *prpE*) genes (92) were individually categorized into the DMSP catabolic sub-category of organic biosynthesis category in CCycDB. DMSP catabolic pathway to release dimethyl sulfide (DMS), which can influence climate and is referred as “climate-active” gas, was listed in the DMS synthesis sub-category of DMSP catabolic pathway. The remaining gene families for organic sulfur transformation and for linking organic and inorganic sulfur transformation processes were included in the organic sulfur cycle sub-category of organic degradation. Inorganic sulfur transformation pathways, such as assimilatory sulfate reduction, dissimilatory sulfur reduction and oxidation, sulfur reduction and oxidation were included in the inorganic sulfur transformation sub-category of organic degradation. Eight classic nitrogen cycle pathways, including N_2_ fixation, anammox, nitrification, denitrification, assimilatory and dissimilatory nitrate reduction, organic nitrogen synthesis and degradation, and hydroxylamine reduction can be found in nitrogen cycle sub-category of organic degradation category. Organic nitrogen compounds (e.g., amino acid and nucleotide) were included in organic biosynthesis and organic degradation category. Key gene families that mediate inorganic and organic phosphorus compound metabolic processes were included in organic biosynthesis, respectively. The sub-category of inorganic phosphorus comprises genes involved in transport inorganic phosphorus compounds (e.g., orthophosphate) outside the member into the cell. The sub-category of organophosphorus compound included cellular phosphorus metabolic processes for synthesizing organic phosphorus compounds, such as pyruvate, phosphonate and phosphinate, purine, and pyrimidine. By generating comprehensive microbial profiles targeting these gene families, the coupling mechanisms among different cycles can be investigated via various statistical approaches.

We applied CCycDB to analyze microbial C cycling gene families in seven different habitats. The functional and taxonomic profiles of C cycling gene families were significantly different among different environments. For CO_2_ fixation, genes encoding ribulose 1,5-bisphosphate carboxylase/oxygenase (RubisCO) were abundant in all tested samples, consistently with our current knowledge that RuBisCO is one of the oldest enzymes on Earth and widely distributed in all ecosystems (93, 94). In contrast, the key enzymes responsible for the ATP-dependent cleavage of citrate to acetyl-CoA and oxaloacetate in the rTCA were rarely detected in these samples, regardless of by ATP citrate lyase or the combined action of citryl-CoA synthetase and citryl-CoA lyase. This observation can be attributed to the more ATP consumption required by the rTCA cycle, which is usually found in anaerobic and sulfur-rich stringent environments (95–97). The key gene *hcd* encoding 4-hydroxybutyryl-CoA dehydratase (98) in the 3HP/4HB and DC/4HB cycles was almost undetectable. So far, this enzyme has been only found to function in few anaerobic bacteria that ferment 4-aminobutyrate (99). Gene families related to CH_4_ fixation and release were most abundant in the mangrove samples, with a higher number of genes detected for aerobic oxidation of methane than anaerobic oxidation. The most abundant pathways of methanogenesis were central methanogenic pathway, hydrogenotrophic and aceticlastic methanogenesis, which is consistent with previous studies (100, 101). Gene families associated with the methane cycle were predominantly present in Proteobacteria, while methanotrophs were mainly distributed in Alphaproteobacteria and Gammaproteobacteria, such as *Methylobacterium*, *Methylocystis*, *Methylococcaceae* (102, 103).

## Conclusions

CCycBD serves as a curated functional gene database for the accurate fingerprinting microbial communities involved in carbon cycling processes. By integrating multiple databases and introducing homologous gene families, CCycDB achieves high performance in terms of coverage, accuracy and specificity. In addition to its role as a reference database, CCycDB functions as a customized knowledge base that enable users to efficiently align sequences and query associated functional information. Compared with existing tools, CCycDB places strong emphasis on usability and accessibility. The workflow simplified the pre-processing steps for input data, and support multiple analysis strategies, including both “read-based” and “assembly-based” approaches. We aim to provide more available information through one-step database searching and integrate multiple annotation from KEGG and CAZy as output, facilitating comprehensive interpretation of the results. Currently, CCycDB contains 4,572 functional genes represented by 10,548,067 curated reference sequences. Extensive validation analyses have demonstrated the high coverage, specificity, and accuracy of CCycDB, highlighting it as a powerful tool for exploring the complex links between microbial carbon cycling processes and environmental change.

## Abbreviations

3-HP: 3-hydroxypropionate bicycle
3-HP/4-HB: 3-hydroxypropionate/4hydroxybutyrate
AAs: Auxiliary activities
BRENDA: Braunschweig enzyme database
C: Carbon
CARD: Comprehensive antibiotic resistance database
CAZy: Carbohydrate-Active Enzymes database
CBB: Calvin-Benson-Bassham
CEs: Carbohydrate esterases
CH_4_: Methane
CO: Carbon monoxide
CO_2_: Carbon dioxide
CODH: Carbon monoxide dehydrogenase
COG: Clusters of Orthologous Groups
DC/4-HB: Dicarboxylate/4-hydroxybutyrate cycles
DCM: Deep chlorophyll maximum layer
DMS: Dimethyl sulfide
DMSP: Dimethylsulphoniopropionate
DRAM: Distilled and refined annotation of metabolism
eggNOG: Evolutionary genealogy of genes -- Non-supervised Orthologous Groups
GHs: Glycoside hydrolases
KEGG: Kyoto Encyclopedia of Genes and Genomes
MCP: Microbial carbon pump
MES: Mesopelagic zone
PCoA: Principal coordinates analysis
PLs: Polysaccharide lyases
RDOC: Recalcitrant dissolved organic carbon
rTCA: Reductive tricarboxylic acid cycle
RuBisCO: Ribulose-1,5-bisphosphate carboxylase/oxygenase
SRF: Surface water layer
WL: Wood-Ljungdahl.

## Data Availability

CCycDB and utilities are available at https://ccycdb.github.io and the ZENODO website under the record number 10045943.

## Code Availability

This study describes the data analysis methods, software, and relevant parameters used in the Methods section. More detailed information is provided in https://github.com/ccycdb/ccycdb.github.io and https://ccycdb.github.io/css/html/usage.html. Where specific parameters are not selected, default parameters were used. The custom scripts are available at https://github.com/ccycdb/ccycdb.github.io.

## Acknowledgements

This study contributes to the science plan of the Ocean Negative Carbon Emissions (ONCE) program. The study was supported by National Key Research and Development Program of China (2020YFA0607600), by the National Natural Science Foundation of China (32371598, 42225708), by the Southern Marine Science and Engineering Guangdong Laboratory (Zhuhai) (SML2024SP022, SML2024SP002), by the Taishan Young Scholarship of Shandong Province, and by the Distinguished Scholarship of Shandong University. The funders had no role in study design, data collection and interpretation, or the decision to submit the work for publication.

## Author Contributions

Qichao Tu conceived the research. Jiayin Zhou and Qichao Tu performed the bioinformatics analyses and wrote the manuscript. Lu Qian and Mengzhi Ji helped with the data analysis. Yan Li, Kai Ma, Xiaoli Yu, Jiyu Chen, Lu Lin, Xiaofan Gong, Zhili He, Jianjun Wang revised the manuscript.

## Ethics statement

Not applicable.

