## Supplementary Figure for "CCycDB: an integrative knowledgebase to fingerprint microbially mediated carbon cycling processes"

**Supplementary materials**

**A.	Supplementary Tables**

**Table S1** Summary of categories and sub-categories in the CCycDB.

**Table S2** Detailed information of gene families included in the CCycDB.

**Table S3** List of CAZy identifiers related to carbohydrate degradation collected from CAZy.

**Table S4** Summary of mapped sequences in the artificial dataset. Red denote misassigned or not well annotated.

**Table S5** Detection rate of related gene families included in the artificial data set at 30% sequence identity.

**Table S6** Detection rate of unrelated gene families included in the artificial dataset at 30% sequence identity.

**Table S7** Summary of shotgun metagenome sequencing data used in this study.

**B.	Supplementary Figures**

**Figure S1** Summary of microbial taxonomic groups associated with carbon cycling processes in CCycDB.

**Figure S2** The composition and relative abundance of microbial gene families (A) and taxonomic groups at family level (B) in seven habitats. For the gene families, the top 20 most abundant gene families were shown. For the taxonomic groups, the top 22 most abundant microbial families were shown. SRF, surface water layer; DCM, deep chlorophyll maximum layer; MES, mesopelagic zone.

**Figure S3** Community dissimilarity of seven habitats based on C cycling gene families (A) and taxonomic groups (species level) (B). The Bray-Curtis dissimilarity considering the relative abundances of gene families and microbial species was used here.

**Figure S4** The distribution of the relative abundance of carbon release gene families in different environments. The numbers above each bar refer to the number of detected gene families involved in carbon cycling processes.

**Figure S5** The relative abundance of predominant microbial taxonomic groups involved in carbon release in different environments.

**B.	Supplementary Figures**

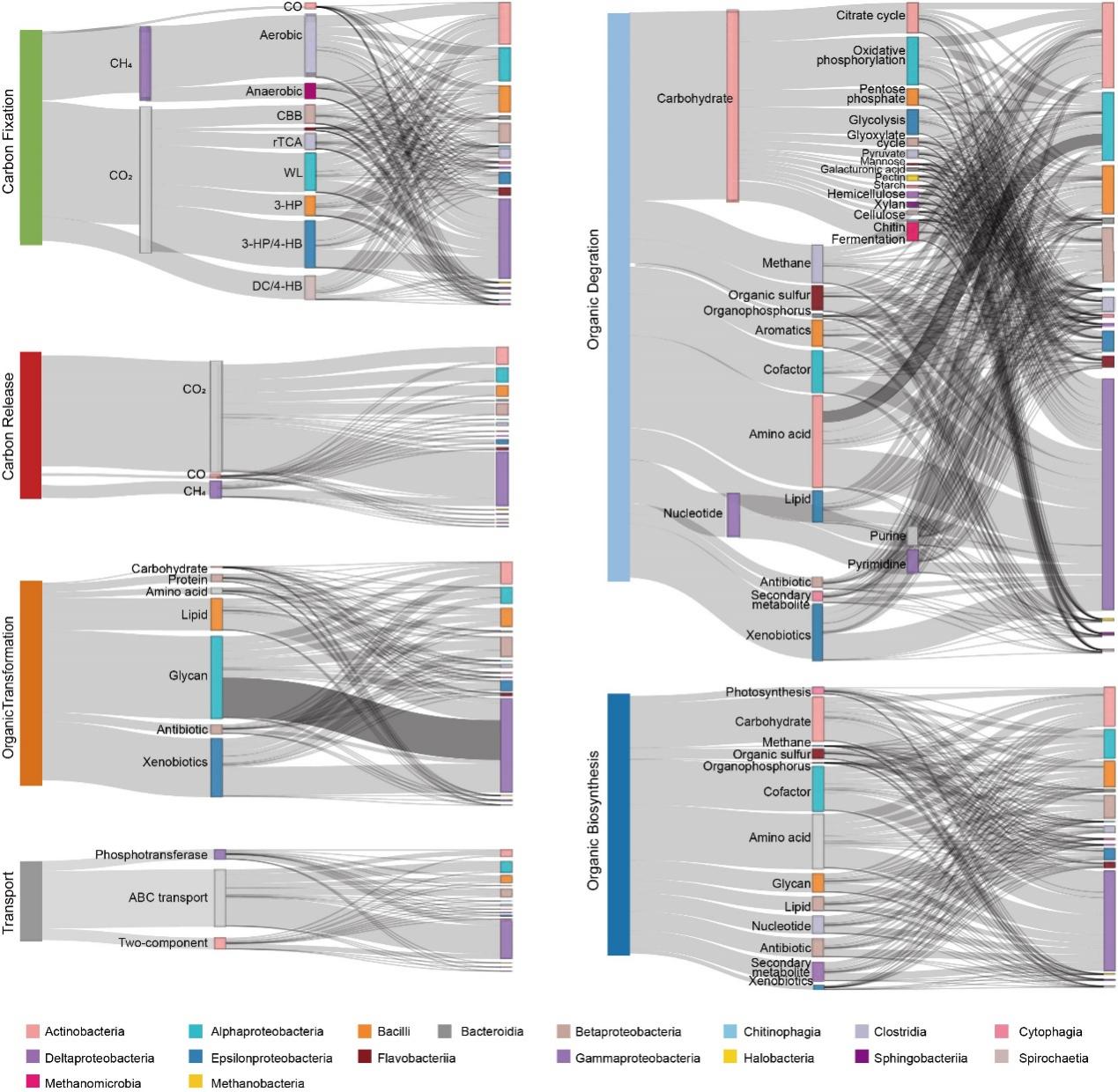

**Figure S1.** Summary of microbial taxonomic groups associated with carbon cycling processes in CCycDB.

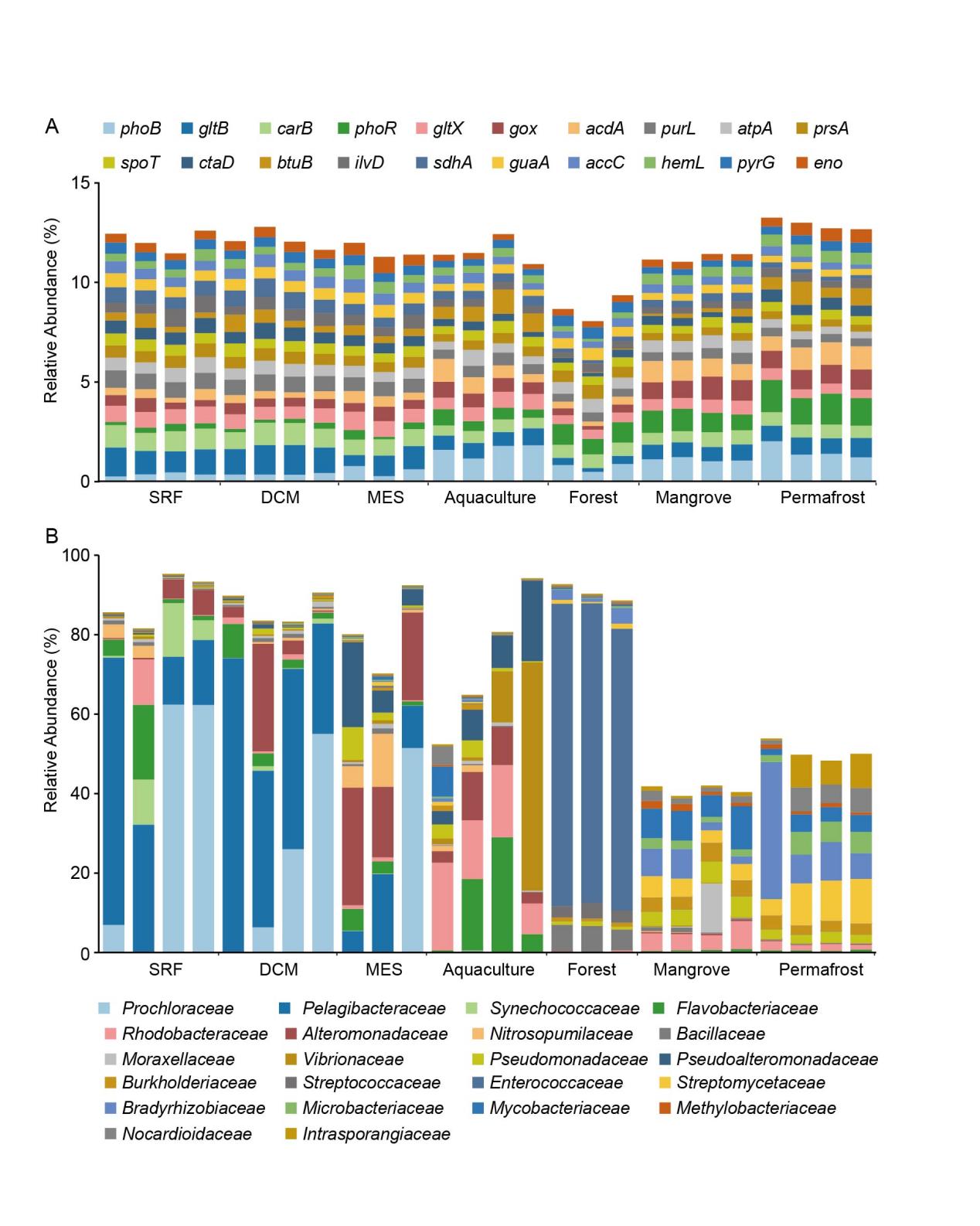

**Figure S2.** The composition and relative abundance of microbial gene families (A) and taxonomic groups at family level (B) in seven habitats. For the gene families, the top 20 most abundant gene families were shown. For the taxonomic groups, the top 22 most abundant microbial families were shown. SRF, surface water layer; DCM, deep chlorophyll maximum layer; MES, mesopelagic zone.

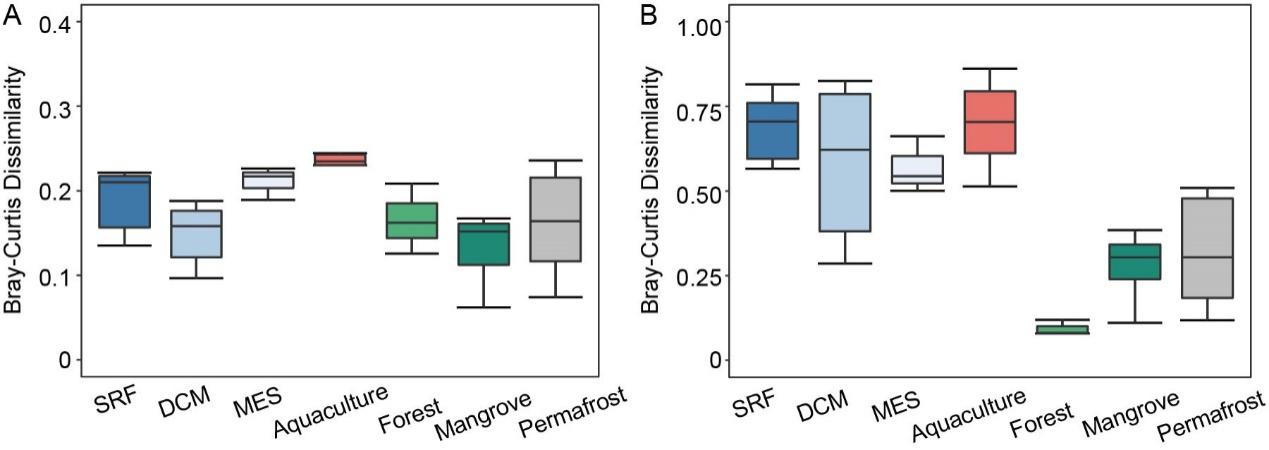

**Figure S3.** Community dissimilarity of seven habitats based on C cycling gene families (A) and taxonomic groups (species level) (B). The Bray-Curtis dissimilarity considering the relative abundances of gene families and microbial species was used here.

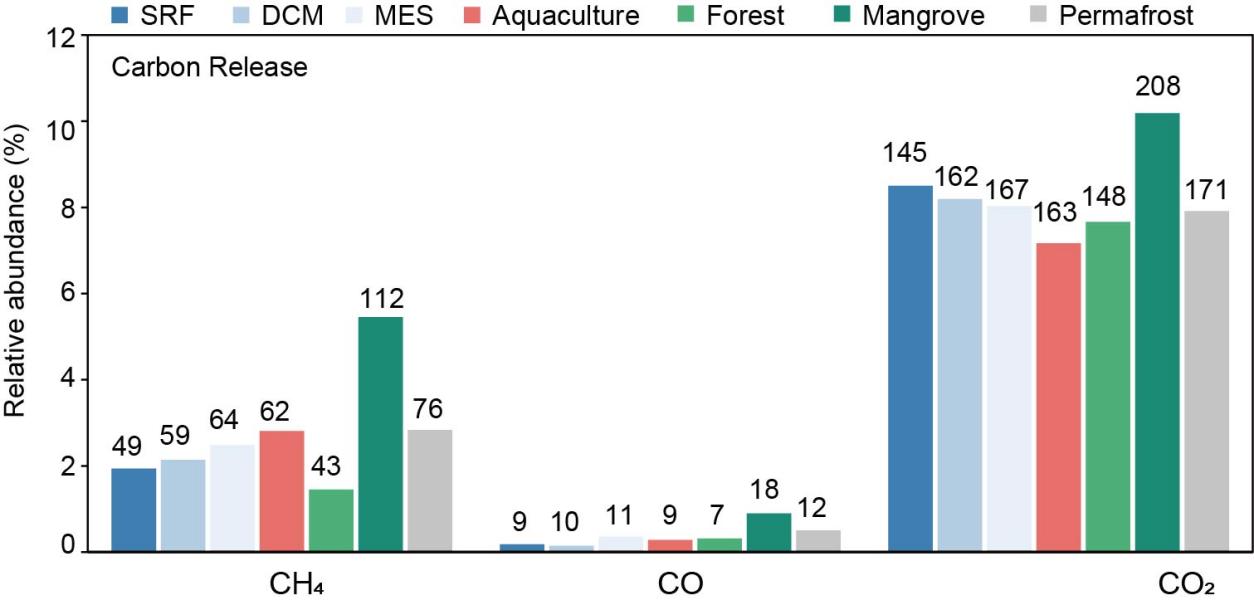

**Figure S4.** The distribution of the relative abundance of carbon release gene families in different habitats. The numbers above each bar refer to the number of detected gene families involved in carbon cycling processes.

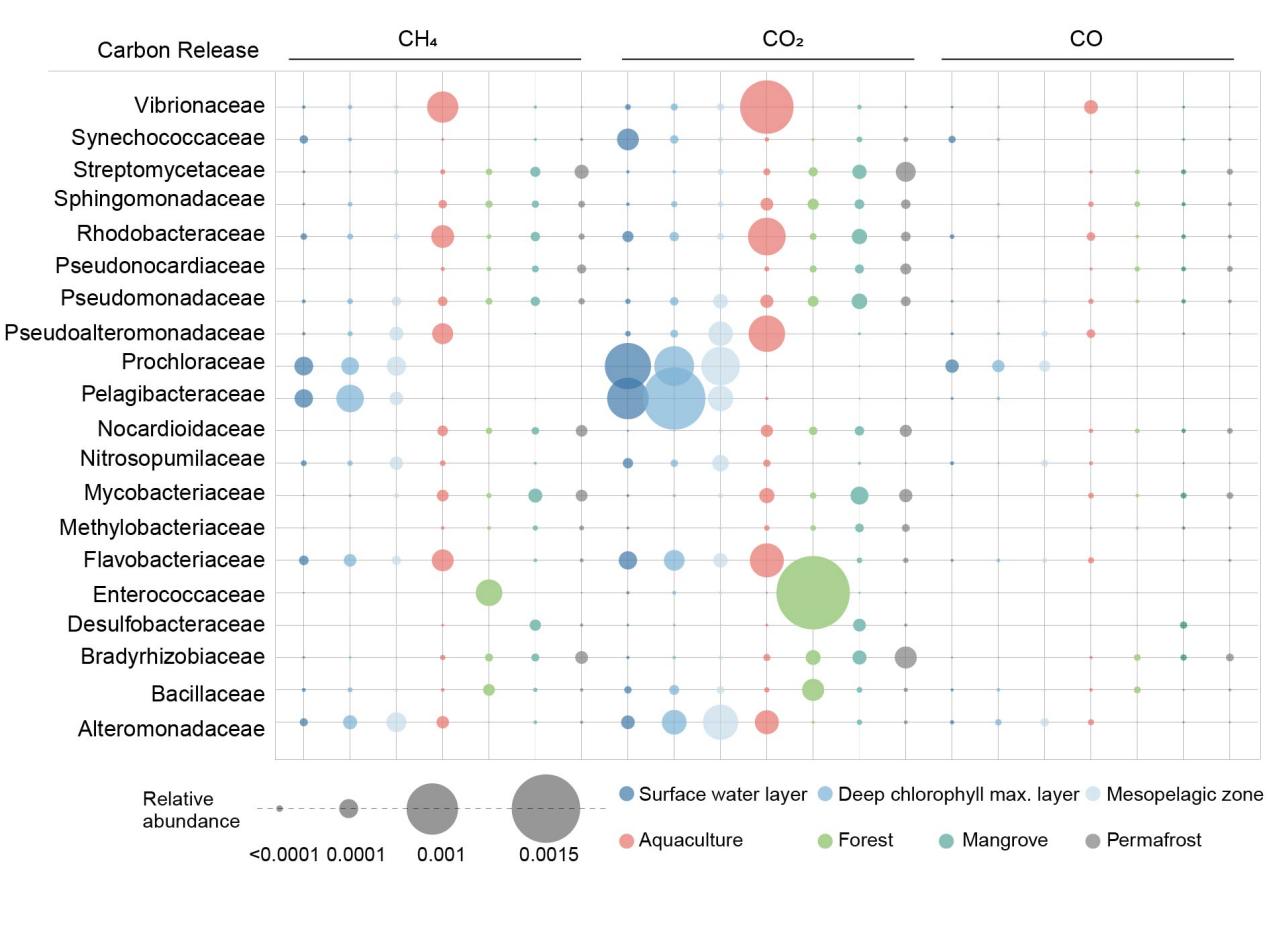

**Figure S5.** The relative abundance of predominant microbial taxonomic groups involved in carbon release in different environments.
