## Supplementary material for "CCycDB: an integrative knowledgebase to fingerprint microbially mediated carbon cycling processes": Table

**Table 1** Statistics of carbon metabolism gene families in CCycDB. Information including gene family numbers, number of sequences, and orthology sequences were summarized.

| **Category** | **SubCategoryI** | **SubCategoryII** | **Gene family** | **Full database** | **Orthology groups with homologous sequences** | | | |
| --- | --- | --- | --- | --- | --- | --- | --- | --- |
|  |  |  |  |  | arCOG | COG | eggNOG | KEGG |
| Carbon Fixation | CH4 |  | 207 | 505,965 | 4,798 | 7,792 | 68,820 | 46,040 |
|  | CO |  | 16 | 8,092 | 91 | 337 | 1,868 | 1,861 |
|  | CO2 | Calvin-Benson-Bassham Cycle | 28 | 119,315 | 8,740 | 3,105 | 25,485 | 18,840 |
|  |  | Reductive tricarboxylic acid cycle | 14 | 12,242 | 469 | 8,054 | 2,846 | 6,950 |
|  |  | Reductive acetyl-CoA pathway | 25 | 110,897 | 784 | 3,090 | 4,677 | 9,147 |
|  |  | 3-Hydroxypropionate Bicycle | 30 | 184,500 | 31 | 1,633 | 11,399 | 9,753 |
|  |  | 3-Hydroxypropionate/4-Hydroxybutyrate Cycle | 20 | 119,652 | 4,553 | 5,968 | 10,479 | 9,132 |
|  |  | Dicarboxylate/4-hydroxybutyrate cycle | 33 | 189,056 | 4,780 | 6,065 | 11,807 | 8,802 |
| Carbon Release | CH4 | Central methanogenic pathway | 121 | 64,653 | 796 | 781 | 6,354 | 7,115 |
|  |  | Aceticlastic methanogenesis | 21 | 76,216 | 237 | 918 | 8,056 | 5,835 |
|  |  | Hydrogenotrophic methanogenesis | 22 | 11,519 | 41 | 63 | 552 | 130 |
|  |  | Methylotrophic methanogenesis | 17 | 9,102 | 3 | 6 | 137 | 864 |
|  | CO |  | 23 | 41,449 | 91 | 574 | 3,582 | 4,264 |
|  | CO2 |  | 337 | 889,450 | 3,631 | 21,191 | 112,815 | 104,035 |
| Organic Biosynthesis | Amino acid |  | 385 | 2,137,146 | 901 | 27,583 | 208,982 | 186,334 |
|  | Antibiotic |  | 666 | 669,509 | 485 | 15,024 | 116,662 | 78,116 |
|  | Carbohydrate |  | 542 | 1,460,277 | 2,944 | 28,901 | 207,843 | 188,254 |
|  | Cofactor |  | 371 | 1,856,467 | 1,595 | 32,472 | 237,842 | 182,282 |
|  | Glycan |  | 301 | 656,645 | 377 | 16,162 | 105,459 | 80,365 |
|  | Lipid |  | 311 | 574,776 | 325 | 10,241 | 104,750 | 64,326 |
|  | Nucleotide |  | 113 | 1,027,165 | 1,448 | 25,517 | 173,542 | 145,044 |
|  | Photosynthesis |  | 69 | 420,838 | 8,819 | 4,688 | 44,256 | 35,510 |
|  | Organophosphorus compound | Organic phosphoester hydrolysis | 13 | 18,404 | 18 | 1,014 | 8,034 | 6,101 |
|  |  | Pentose phosphate pathway | 8 | 49,974 | 371 | 1,892 | 12,392 | 11,428 |
|  |  | Phosphonate and phosphinate metabolism | 34 | 33,214 | 4 | 854 | 5,981 | 5,038 |
|  |  | Phosphotransferase system | 2 | 11,192 | 16 | 763 | 5,539 | 3,323 |
|  |  | Purine metabolism | 25 | 337,859 | 2,460 | 11,596 | 70,188 | 51,678 |
|  |  | Pyrimidine metabolism | 18 | 238,297 | 1,386 | 7,213 | 41,597 | 31,517 |
|  |  | Pyruvate metabolism | 6 | 48,457 | 297 | 1,899 | 12,569 | 10,339 |
|  |  | Transmembrane transport | 18 | 73,580 | 113 | 3,530 | 25,974 | 10,368 |
|  |  | Two-component system | 3 | 670 | 0 | 16 | 75 | 78 |
|  | Inorganic  Phosphorus | Oxidative phosphorylation | 2 | 32,190 | 157 | 1,044 | 5,713 | 4,512 |
|  |  | Transmembrane transport | 10 | 52,475 | 492 | 2,845 | 15,888 | 11,752 |
|  |  | Two-component system | 6 | 50,196 | 9 | 5,378 | 33,144 | 5,263 |
|  |  | Auxiliary | 6 | 7,520 | 0 | 660 | 3,987 | 2,561 |
|  | Secondary metabolite |  | 547 | 665,574 | 465 | 15,066 | 113,854 | 81,259 |
|  | Xenobiotics |  | 24 | 158,231 | 58 | 1,586 | 11,274 | 11,595 |
|  | Amino acid |  | 385 | 2,137,146 | 901 | 27,583 | 208,982 | 186,334 |
|  | Others |  | 144 | 171516 | 80 | 4738 | 34107 | 17470 |
| Organic Degration | Amino acid |  | 310 | 1,143,709 | 5,240 | 25,106 | 182,665 | 168,521 |
|  | Antibiotic |  | 56 | 156,044 | 141 | 4,221 | 22,167 | 22,930 |
|  | Aromatics |  | 350 | 291,060 | 2,009 | 7,256 | 63,343 | 41,901 |
|  | Carbohydrate | Citrate cycle | 39 | 254,875 | 415 | 2,291 | 15,383 | 15,410 |
|  |  | Oxidative phosphorylation | 112 | 860,722 | 397 | 4,760 | 33,143 | 31,662 |
|  |  | Pentose phosphate | 67 | 227,115 | 565 | 4,444 | 39,670 | 31,951 |
|  |  | Glyoxylate cycle | 5 | 87,576 | 111 | 762 | 5,686 | 6,074 |
|  |  | Glycolysis | 68 | 303,539 | 728 | 6,664 | 60,057 | 46,409 |
|  |  | Pyruvate | 26 | 97,321 | 362 | 3,156 | 19,731 | 24,814 |
|  |  | Mannose | 2 | 15,172 | 1 | 85 | 2,133 | 548 |
|  |  | Galactose | 11 | 5,631 | 1 | 1,016 | 8,414 | 9,438 |
|  |  | Arabinose | 16 | 29,907 | 247 | 743 | 10,186 | 7,174 |
|  |  | Xylose | 6 | 22,865 | 0 | 299 | 3,346 | 3,359 |
|  |  | Fucose | 5 | 7,046 | 1 | 14 | 195 | 106 |
|  |  | Galacturonic acid | 3 | 26,163 | 1 | 64 | 454 | 340 |
|  |  | Starch | 45 | 60,444 | 13,935 | 6,547 | 15,566 | 16,400 |
|  |  | Hemicellulose | 55 | 65,450 | 103 | 90,905 | 18,110 | 17,224 |
|  |  | Mannan | 25 | 33,774 | 250 | 2,425 | 12,957 | 10,090 |
|  |  | Arabinan | 21 | 30,268 | 16 | 88,461 | 10,240 | 9,283 |
|  |  | Xylan | 36 | 56,649 | 18 | 2,803 | 13,334 | 13,082 |
|  |  | Cellulose | 32 | 42,449 | 254 | 2,791 | 17,173 | 14,761 |
|  |  | Pectin | 38 | 59,258 | 250 | 1,306 | 13,893 | 10,379 |
|  |  | Chitin | 32 | 33,417 | 20 | 3,248 | 101,350 | 11,539 |
|  |  | Fermentation | 34 | 190,083 | 219 | 1,954 | 15,317 | 12,472 |
|  |  | Others | 480 | 883,036 | 3,226 | 116,583 | 198,543 | 156,374 |
|  | Cofactor | Coenzyme | 11 | 58,460 | 28 | 527 | 2,787 | 2,328 |
|  |  | Porphyrin | 6 | 41,760 | 4 | 165 | 496 | 414 |
|  |  | Vitamin A | 4 | 19,504 | 12 | 24 | 2,050 | 1,727 |
|  |  | Vitamin B | 136 | 585,032 | 408 | 16,074 | 104,939 | 77,790 |
|  |  | Vitamin E | 36 | 49,203 | 21 | 1,714 | 10,719 | 9,421 |
|  |  | Vitamin K | 3 | 4,889 | 0 | 59 | 249 | 428 |
|  | Lipid | Fatty acid | 69 | 384,361 | 72 | 3,934 | 33,181 | 31,676 |
|  |  | Polyketide | 7 | 2,858 | 2 | 82 | 888 | 67 |
|  |  | Others | 27 | 24,110 | 806 | 1,296 | 10,975 | 7,575 |
|  | Methane | Oxidation of formaldehyde | 10 | 20,467 | 3,495 | 59 | 649 | 1,154 |
|  |  | Oxidation of formate | 12 | 10,199 | 61 | 103 | 453 | 929 |
|  |  | Oxidation of merthane and C1 compounds | 50 | 80,056 | 219 | 1,534 | 13,053 | 6,627 |
|  |  | RuMP pathway | 15 | 70,137 | 80 | 401 | 8,748 | 4,647 |
|  |  | serine cycle | 29 | 216,755 | 675 | 5,133 | 38,008 | 25,864 |
|  |  | Anaerobic oxidation of methane | 96 | 114,748 | 391 | 1,099 | 8,726 | 8,616 |
|  | Nitrogen cycle | Anammox | 5 | 2,105 | 0 | 1 | 3 | 0 |
|  |  | Assimilatory nitrate reduction | 6 | 8,263 | 0 | 84 | 222 | 1,005 |
|  |  | Auxiliary | 4 | 13,840 | 0 | 12 | 24 | 3 |
|  |  | Denitrification | 16 | 56,618 | 15 | 285 | 1,749 | 916 |
|  |  | Dissimilatory nitrate reduction | 17 | 28,984 | 9 | 170 | 1,097 | 794 |
|  |  | Nitrification | 9 | 38,391 | 0 | 5 | 5 | 1 |
|  |  | Nitrogen fixation | 5 | 18,311 | 0 | 13 | 19 | 25 |
|  |  | Organic degradation and synthesis | 17 | 114,363 | 6 | 971 | 5,131 | 4,978 |
|  | DMSP catabolic pathway | DMS dehydrogenase | 1 | 27 | 0 | 0 | 0 | 0 |
|  |  | DMS monooxgenase | 1 | 1,900 | 2 | 2 | 36 | 9 |
|  |  | DMS synthesis | 6 | 6,617 | 2 | 6 | 39 | 23 |
|  |  | DMSP demethylation | 4 | 6,234 | 2 | 6 | 34 | 21 |
|  |  | DMSP lyase | 7 | 441 | 1 | 1 | 11 | 6 |
|  |  | DMSP synthesis | 2 | 128 | 0 | 0 | 40 | 6 |
|  |  | DMSP trasnport | 1 | 103 | 0 | 0 | 7 | 2 |
|  |  | Ancillary genes | 7 | 11,711 | 9 | 16 | 236 | 68 |
|  | Organic sulfur cycle | Auxiliary | 31 | 112,103 | 200 | 333 | 4,055 | 1,062 |
|  |  | Link between inorganic and organic sulfur transformation | 35 | 144,620 | 136 | 194 | 3,131 | 583 |
|  |  | Organic sulfur transformation | 57 | 147,231 | 144 | 257 | 3,262 | 1,060 |
|  | Inorganic sulfur transformation | Assimilatory sulfate reduction | 11 | 117,455 | 117 | 177 | 3,767 | 519 |
|  |  | Dissimilatory sulfur reduction and oxidation | 22 | 20,354 | 41 | 43 | 573 | 118 |
|  |  | SOX systems | 7 | 14,998 | 14 | 35 | 694 | 108 |
|  |  | Sulfur disproportionation | 5 | 1,394 | 4 | 7 | 39 | 14 |
|  |  | Sulfur oxidation | 14 | 15,372 | 3 | 21 | 161 | 46 |
|  |  | Sulfur reduction | 26 | 11,546 | 46 | 64 | 305 | 83 |
|  | Nucleotide | Purine | 57 | 361,274 | 2,234 | 12,645 | 81,132 | 60,826 |
|  |  | Pyrimidine | 45 | 270,396 | 1,387 | 9,138 | 51,905 | 44,090 |
|  | Xenobiotics | Lignin | 641 | 723287 | 6391 | 26867 | 203029 | 127608 |
|  | Secondary metabolite |  | 203 | 238,301 | 577 | 6,431 | 48,759 | 38,078 |
|  | Others |  | 26 | 46,506 | 98 | 3,090 | 20,071 | 5,414 |
| Organic Transformation | Amino acid |  | 19 | 31,623 | 0 | 1,187 | 7,415 | 8,034 |
|  | Antibiotic |  | 147 | 48,536 | 36 | 939 | 9,154 | 5,277 |
|  | Carbohydrate |  | 8 | 4,259 | 3 | 79 | 915 | 437 |
|  | Glycan |  | 139 | 477,974 | 254 | 8,283 | 56,144 | 45,053 |
|  | Lipid |  | 214 | 172,714 | 199 | 4,850 | 46,997 | 38,088 |
|  | Protein |  | 13 | 64,533 | 9 | 762 | 5,144 | 4,429 |
|  | Xenobiotics |  | 248 | 379,903 | 620 | 13,580 | 107,327 | 70,195 |
| Transport | ABC transport |  | 331 | 952,843 | 970 | 19,012 | 131,020 | 104,293 |
|  | Phosphotransferase system |  | 86 | 148,410 | 585 | 4,966 | 38,166 | 24,456 |
|  | Transport system |  | 6 | 10,233 | 1 | 98 | 3,447 | 2,794 |
|  | Two-component system |  | 87 | 250,348 | 823 | 9,458 | 55,800 | 22,175 |
|  | Others |  | 18 | 61,363 | 18 | 3,532 | 20,297 | 8,632 |

| **Category** | **SubCategory I** | **SubCategory II** | **Pathway** | **Gene families** |
| --- | --- | --- | --- | --- |
| Carbon Fixation | 4 | 17 | 21 | 413 |
| Carbon Release | 3 | 6 | 6 | 499 |
| Organic Biosynthesis | 12 | 55 | 85 | 2,425 |
| Organic Degration | 15 | 91 | 111 | 2,307 |
| Organic Transformation | 7 | 15 | 19 | 720 |
| Transport | 4 | 4 | 4 | 500 |

### Table 2. Summary of gene families and pathways for different categories in CCycDB.

**Table 3** Summary of the microbial taxa at different taxonomic levels for the recruited sequences in CCycDB

|  |  | **Phylum** | | **Class** | | **Order** | | **Family** | | **Genus** | | **Species** | |
| --- | --- | --- | --- | --- | --- | --- | --- | --- | --- | --- | --- | --- | --- |
|  |  | Archaea | Bacteria | Archaea | Bacteria | Archaea | Bacteria | Archaea | Bacteria | Archaea | Bacteria | Archaea | Bacteria |
| Carbon Fixation | CH4-arobic | 5 | 42 | 10 | 71 | 19 | 176 | 32 | 412 | 97 | 2,163 | 350 | 15,231 |
|  | CH4-anaerobic | 5 | 31 | 9 | 53 | 14 | 130 | 24 | 290 | 80 | 1,299 | 266 | 6,412 |
|  | CO | 3 | 20 | 6 | 39 | 12 | 78 | 19 | 131 | 41 | 403 | 99 | 1,452 |
|  | CO2-CBB | 4 | 38 | 7 | 73 | 17 | 177 | 26 | 411 | 63 | 2,107 | 153 | 12,938 |
|  | CO2-rTCA | 4 | 13 | 5 | 17 | 8 | 54 | 11 | 112 | 13 | 354 | 29 | 1,388 |
|  | CO2-WL | 4 | 36 | 8 | 67 | 17 | 172 | 30 | 383 | 89 | 1,892 | 276 | 10,914 |
|  | CO2-3HP | 5 | 39 | 8 | 72 | 15 | 178 | 24 | 410 | 88 | 2,149 | 286 | 14,461 |
|  | CO2-3HP/4HB | 3 | 36 | 6 | 67 | 11 | 172 | 14 | 380 | 29 | 1,933 | 69 | 11,426 |
|  | CO2-DC/4HB | 5 | 36 | 11 | 69 | 21 | 168 | 35 | 396 | 111 | 2,056 | 377 | 14,171 |
|  | Photosynthesis | 3 | 40 | 10 | 70 | 15 | 181 | 22 | 429 | 70 | 2,238 | 176 | 12,825 |
| Carbon Release | CH4 | 5 | 37 | 11 | 67 | 17 | 160 | 27 | 371 | 90 | 1,801 | 314 | 11,735 |
|  | CO | 4 | 25 | 10 | 53 | 18 | 122 | 28 | 271 | 67 | 1,008 | 189 | 5,036 |
|  | CO2 | 5 | 48 | 11 | 82 | 23 | 195 | 41 | 456 | 130 | 2,461 | 504 | 18,367 |
| Organic Biosynthesis | Amino acid | 6 | 53 | 11 | 81 | 23 | 195 | 41 | 460 | 138 | 2,587 | 567 | 19,939 |
|  | Antibiotic | 6 | 51 | 11 | 78 | 21 | 189 | 38 | 444 | 115 | 2,460 | 429 | 18,194 |
|  | Carbohydrate | 6 | 53 | 11 | 80 | 23 | 193 | 39 | 459 | 128 | 2,532 | 499 | 19,431 |
|  | Cofactor | 7 | 54 | 11 | 82 | 23 | 196 | 40 | 460 | 139 | 2,564 | 567 | 19,748 |
|  | Glycan | 5 | 53 | 9 | 78 | 17 | 191 | 28 | 447 | 70 | 2,459 | 182 | 17,936 |
|  | Lipid | 6 | 52 | 11 | 77 | 21 | 189 | 32 | 442 | 105 | 2,411 | 336 | 17,346 |
|  | Methane | 5 | 34 | 11 | 51 | 22 | 120 | 34 | 288 | 106 | 1,044 | 379 | 5,481 |
|  | Nucleotide | 6 | 51 | 11 | 79 | 22 | 191 | 39 | 453 | 125 | 2,471 | 463 | 17,946 |
|  | Organophosphorus compound | 6 | 49 | 11 | 73 | 20 | 178 | 36 | 407 | 109 | 2,227 | 400 | 14,172 |
|  | Inorganic phosphorus | 6 | 42 | 9 | 70 | 16 | 163 | 24 | 362 | 84 | 1,808 | 275 | 8,029 |
|  | Photosynthesis | 5 | 48 | 10 | 76 | 21 | 186 | 33 | 437 | 97 | 2,379 | 271 | 16,165 |
|  | Secondary metabolite | 6 | 51 | 11 | 78 | 21 | 189 | 38 | 444 | 115 | 2,460 | 429 | 18,175 |
|  | Xenobiotics | 5 | 43 | 8 | 68 | 14 | 163 | 19 | 382 | 33 | 1,867 | 80 | 12,528 |
| Organic Degration | Amino acid | 6 | 51 | 11 | 78 | 22 | 188 | 39 | 446 | 125 | 2,450 | 509 | 18,439 |
|  | Antibiotic | 6 | 41 | 10 | 63 | 17 | 161 | 25 | 380 | 60 | 1,852 | 142 | 11,245 |
|  | Aromatics | 6 | 45 | 11 | 72 | 20 | 174 | 31 | 401 | 101 | 2,165 | 310 | 15,044 |
|  | Citrate cycle | 6 | 47 | 11 | 73 | 21 | 181 | 35 | 418 | 115 | 2,279 | 377 | 15,968 |
|  | Oxidative phosphorylation | 6 | 51 | 11 | 79 | 22 | 192 | 36 | 451 | 118 | 2,476 | 434 | 17,630 |
|  | Pentose phosphate | 6 | 45 | 9 | 74 | 19 | 180 | 30 | 415 | 99 | 2,189 | 352 | 14,722 |
|  | Glyoxylate cycle | 5 | 37 | 9 | 62 | 17 | 160 | 28 | 377 | 81 | 1,965 | 250 | 12,669 |
|  | Glycolysis | 6 | 49 | 10 | 75 | 20 | 184 | 35 | 422 | 108 | 2,250 | 372 | 15,842 |
|  | Pyruvate | 4 | 35 | 9 | 60 | 13 | 152 | 22 | 344 | 52 | 1,707 | 136 | 11,702 |
|  | Mannose | 1 | 18 | 1 | 32 | 1 | 72 | 1 | 150 | 1 | 525 | 3 | 3,107 |
|  | Galactose | 1 | 17 | 1 | 26 | 1 | 52 | 1 | 101 | 1 | 264 | 2 | 1,221 |
|  | Xylose | 1 | 21 | 2 | 34 | 4 | 82 | 7 | 178 | 46 | 669 | 196 | 3,752 |
|  | Fucose | 1 | 18 | 2 | 27 | 2 | 50 | 3 | 105 | 4 | 297 | 14 | 1,034 |
|  | Galacturonic acid | 1 | 26 | 1 | 43 | 3 | 108 | 6 | 230 | 38 | 1,147 | 152 | 6,692 |
|  | Starch | 0 | 21 | 0 | 32 | 0 | 75 | 0 | 155 | 0 | 525 | 0 | 3,307 |
|  | Hemicellulose | 1 | 29 | 2 | 51 | 2 | 125 | 2 | 280 | 1 | 1,162 | 4 | 7,431 |
|  | Xylan | 1 | 29 | 2 | 49 | 2 | 119 | 2 | 266 | 1 | 1,113 | 4 | 7,097 |
|  | Cellulose | 0 | 10 | 1 | 38 | 2 | 97 | 2 | 223 | 1 | 862 | 3 | 4,810 |
|  | Pectin | 2 | 29 | 2 | 49 | 2 | 122 | 2 | 270 | 2 | 1,142 | 5 | 7,374 |
|  | Chitin | 1 | 23 | 1 | 36 | 2 | 90 | 2 | 202 | 1 | 780 | 3 | 4,146 |
|  | Fermentation | 6 | 43 | 9 | 70 | 18 | 179 | 29 | 411 | 97 | 2,213 | 304 | 15,169 |
|  | Cofactor | 6 | 51 | 11 | 78 | 22 | 190 | 36 | 449 | 119 | 2,443 | 427 | 17,611 |
|  | Lipid | 4 | 49 | 8 | 77 | 14 | 189 | 18 | 441 | 58 | 2,394 | 170 | 16,645 |
|  | serine cycle | 6 | 47 | 10 | 72 | 20 | 177 | 34 | 419 | 104 | 2,231 | 381 | 15,999 |
|  | Purine | 6 | 46 | 11 | 74 | 22 | 183 | 37 | 432 | 113 | 2,276 | 394 | 15,249 |
|  | Pyrimidine | 6 | 51 | 11 | 78 | 20 | 187 | 34 | 436 | 104 | 2,338 | 353 | 15,865 |
|  | DMSP catabolic pathway | 0 | 11 | 0 | 14 | 0 | 49 | 0 | 99 | 0 | 381 | 0 | 1,891 |
|  | Organic sulfur cycle | 5 | 52 | 10 | 81 | 17 | 183 | 26 | 412 | 57 | 2,197 | 177 | 15,400 |
|  | Inorganic sulfur transformation | 6 | 45 | 8 | 77 | 17 | 171 | 28 | 390 | 64 | 1,896 | 157 | 12,022 |
|  | Secondary metabolite | 6 | 41 | 10 | 63 | 17 | 161 | 25 | 381 | 61 | 1,861 | 143 | 11,376 |
|  | Lignin | 6 | 49 | 11 | 76 | 21 | 184 | 36 | 434 | 116 | 2,297 | 412 | 16,633 |
| Organic Transformation | Amino acid | 1 | 24 | 2 | 38 | 4 | 96 | 5 | 226 | 13 | 915 | 21 | 4,823 |
|  | Antibiotic | 5 | 35 | 10 | 50 | 17 | 129 | 25 | 286 | 43 | 1,173 | 81 | 6,588 |
|  | Carbohydrate | 0 | 11 | 0 | 17 | 0 | 36 | 0 | 71 | 0 | 171 | 0 | 644 |
|  | Glycan | 4 | 52 | 9 | 77 | 13 | 185 | 18 | 436 | 29 | 2,367 | 48 | 16,559 |
|  | Lipid | 5 | 46 | 10 | 68 | 16 | 170 | 25 | 390 | 85 | 1,946 | 251 | 12,734 |
|  | Protein | 6 | 43 | 9 | 61 | 17 | 144 | 23 | 322 | 61 | 1,337 | 129 | 6,605 |
|  | Xenobiotics | 6 | 47 | 11 | 75 | 21 | 184 | 37 | 432 | 116 | 2,250 | 393 | 15,577 |
| Transport | ABC transport | 6 | 50 | 11 | 79 | 20 | 190 | 33 | 445 | 102 | 2,454 | 392 | 18,397 |
|  | Phosphotransferase system | 5 | 46 | 11 | 75 | 19 | 173 | 29 | 379 | 79 | 1,963 | 270 | 13,028 |
|  | Two-component system | 6 | 47 | 11 | 74 | 19 | 177 | 31 | 403 | 99 | 2,163 | 317 | 14,663 |
